# Multiplexed chromatin imaging reveals predominantly pairwise long-range coordination between *Drosophila* Polycomb genes

**DOI:** 10.1101/2022.05.16.492046

**Authors:** Julian Gurgo, Jean-Charles Walter, Jean-Bernard Fiche, Christophe Houbron, Marie Schaeffer, Giacomo Cavalli, Frédéric Bantignies, Marcelo Nollmann

## Abstract

Polycomb (Pc) group proteins are transcriptional regulators with key roles in development, cell identity and differentiation. Pc-bound chromatin regions form repressive domains that interact in 3D to assemble repressive nuclear compartments. Here, we used multiplexed chromatin imaging to investigate whether Pc compartments involve the clustering of multiple Pc domains during Drosophila development. Notably, 3D proximity between Pc targets is rare and involves predominantly pairwise interactions. These 3D proximities are particularly enhanced in segments where Pc genes are co-repressed. In addition, segment-specific expression of *Hox* Pc targets leads to their spatial segregation from Pc repressed genes. Finally, non-Hox Pc targets are proximal in regions where they are co-expressed. These results indicate that long-range Pc interactions are temporally and spatially regulated during differentiation and development but do not involve clustering of multiple distant Pc genes.

## Introduction

Chromatin is folded in a hierarchical, multi-level organization, ranging from nucleosomes to chromosome territories. Chromosome conformation capture techniques revealed that at an intermediate level, chromatin is organized in Topologically Associating Domains (TADs) (Dixon et al. 2012; Nora et al. 2012; Sexton et al. 2012; Hou et al. 2012). TADs can affect several biological processes, like gene regulation (Cavalli and Misteli 2013; Andrey and Mundlos 2017). In *Drosophila*, TADs are generally smaller than in mammals (median size of ~ 100 kb versus ~ 800 kb, respectively) and tend to correlate with active and repressed chromatin marks (Sexton et al. 2012; Szabo, Bantignies, and Cavalli 2019). *Drosophila* TADs are classified according to their epigenetic state into active (associated with H3K4me3 and H3K36me3), heterochromatic (enriched in H3K9me2, HP1, Su(var)), black (devoid of specific histone marks) and Polycomb (Pc) repressed, enriched in H3K27me3 and bound by Pc group (PcG) proteins.

PcG proteins are conserved in most eukaryotic organisms. They mediate gene repression, and are classified into two main complexes, PRC1 and PRC2. In *Drosophila*, PcG proteins are recruited to specific sequences called Polycomb Response Elements (PREs) (Nègre et al. 2006) by a two-step process (Wang et al. 2004) involving the deposition of H3K27me3 marks by PRC2, followed by chromatin compaction by PRC1 (Francis, Kingston, and Woodcock 2004; Grau et al. 2011), a complex that contains the chromodomain protein Polycomb (PC), and Polyhomeotic (PH). PcG target genes are often contained within discrete Pc TADs (Sexton, 2012), which are characterized by a high degree of compaction and intermixing (Boettiger et al. 2016; Wani et al. 2016). In single cells, Pc TADs form discrete nano-compartments (Szabo et al. 2018) displaying a cell-type specific internal organization that responds to the transcriptional state of its PcG genes (Mateo et al. 2019). Interestingly, PcG proteins form discrete compartments (foci) in the nucleus, both in flies and mammals (Saurin et al. 1998; Buchenau et al. 1998; Bantignies et al. 2011). Within these Pc foci, genomically-distant PcG target genes can physically colocalize when co-repressed (Lanzuolo et al. 2007; Bantignies et al. 2011). Notably, recent studies showed that mammalian PRC1 components can form phase-separated compartments (Plys et al. 2019), suggesting that PcG genes within Pc TADs may coalesce in 3D to form phase-separated condensates.

In Drosophila, Pc TADs tend to be spatially segregated from active domains (Boettiger et al. 2016; Szabo et al. 2018), consistent with the formation of active and repressive compartments (Sexton et al. 2012), and with the spatial separation of active and repressive marks in single cells (Boettiger et al. 2016; Cattoni et al. 2017). Taken together, this evidence suggests the possibility that multiple Pc TADs might often associate with each other in single cells to form Pc foci to reinforce gene repression.

Here, we tested this hypothesis by applying Hi-M, an imaging-based method that enables the capture of chromatin conformations in single cells while preserving the spatial information within the specimen (Cardozo Gizzi et al. 2019; 2020). We found that at the chromosomal scale the organization of Pc domains can be well described by a weakly microphase-separated polymer. In addition, spatial colocalization of PcG genes into hubs was rare, and in most cases involved only two Pc domains. Formation of hubs involving more than two Pc domains was highly infrequent. Interestingly, the interaction frequencies between Pc domains were enhanced in regions of the embryo where PcG target genes were co-repressed or co-expressed, indicating that the rare 3D encounters between Pc domains may play a role at reinforcing repression and in co-transcriptional activation.

## Results

### Chromosome-wide, simultaneous visualization of multiple Polycomb targets in single cells

To investigate the chromosome-wide organization of Pc target genes, we focused on a ^~^15Mb region of chromosome 3R (chr3R) displaying most long-range Pc contacts (Sexton et al. 2012; Bantignies et al. 2011; Tolhuis et al. 2011; Hug et al. 2017; Ing-Simmons et al. 2021; Loubiere et al. 2020). Within this region, we identified 19 loci displaying binding of all PRC1 components (PC, PH and PSC) (Fig. 1A and S1A), and involved in most long-range Pc interactions in this chromosome (>80%, Fig. S1B). This selection included PcG targets within the three large Pc TADs containing *Drosophila’s hox* genes: Bithorax (BX-C), Antennapedia (ANT-C) and NK-C, in addition to 8 smaller non-hox PcG-regulated target loci (Fig. 1A). Next, we used Hi-M, an imaging-based technology that we recently developed to retrieve chromatin architecture in single cells while maintaining spatial context (Cardozo Gizzi et al. 2019; 2020). We note that Hi-M is similar to other multiplexed imaging technologies developed concurrently (Mateo et al. 2019; Bintu et al. 2018; Takei et al. 2021; Su et al. 2020; Nir et al. 2018; Liu et al. 2020). Hi-M relies on the sequential imaging of tens of distinct genomic loci labeled by oligopaint-FISH (Beliveau et al. 2015; 2012) in intact *Drosophila* embryos (Cardozo Gizzi et al. 2019, 2020) (Figs. 1B, and S2A-B). Multiplexed DNA imaging methods, such as Hi-M, were previously used in samples with single cell layers (e.g. cultured-cells, *Drosophila* embryos before gastrulation, or cryo-sections). Here, we coupled Hi-M to confocal imaging to be able to visualize the 3D localization of multiple Pc target regions in multi-layered stage 15 - 16 (S15-S16, 12-16h of development) embryos without cryo-dissection (Fig. 1B).

**Figure 1:**
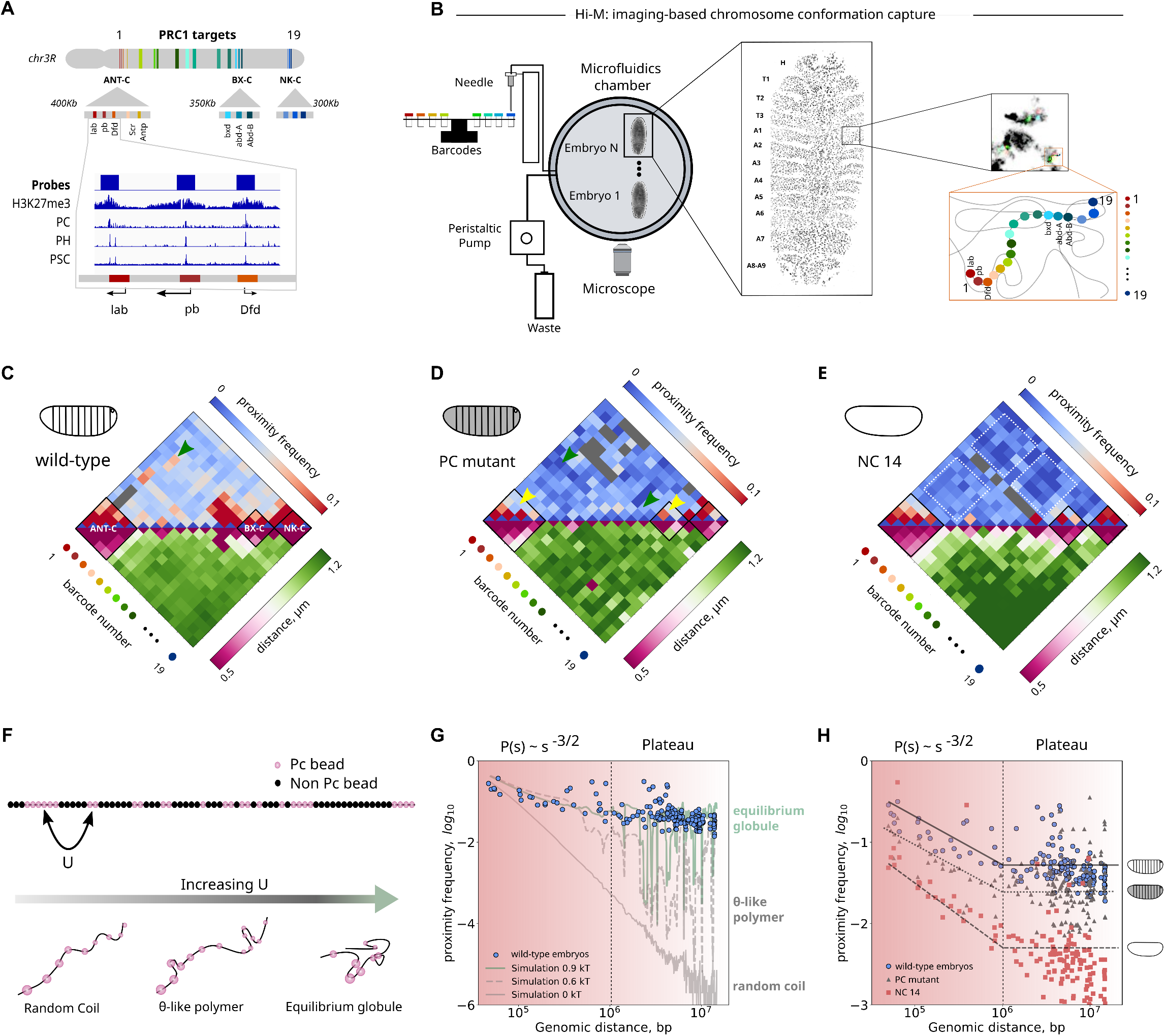
Long-range proximities between distant Polycomb domains are best described by a self-interacting polymer in the globule regime. **A.** Schematic representation of the Oligopaints library used, covering Pc domains over a portion of ^~^ 12Mb of chromosome 3R. Triangles represent the three larger domains: ANTP-C, BX-C and NK-C. Bottom: H3K27me3, PC, PH, PSC Chip-seq data for a region of the ANT-C domain. **B.** Diagram of the experimental setup. Hi-M allows to image Pc domains sequentially in single cells of full embryos, keeping spatial resolution. Briefly, images are acquired using an Airyscan confocal microscope coupled to a fluidics system. Barcodes are injected using a needle attached to a translation stage. A peristaltic pump delivers solutions into the microfluidics chamber. See supplementary data for a detailed description of the setup. **C - E.** Hi-M proximity frequency maps (top) and pairwise distance maps (bottom) for stage 15-16 embryos (C, N=22243, n=20, 5 replicates), Pc mutant (D, N=9169, n=7, 1 replicate), and NC14 wild-type embryos (E, N=12625, n=10, 1 replicate). Barcode IDs are indicated on the axis. Colormaps corresponding to the proximity frequency and pairwise distance are shown above and below the maps, respectively. Gray bins in the matrix correspond to interactions that did not satisfy our quality control filters (Methods). Proximity frequency maps were calculated with a distance threshold of 250 nm. **F.** Polymer modeling of chromosome 3R. Co-polymer containing interacting beads with energy *U* (Pc beads, pink), and non-interacting beads (black). Increasing *U* leads to three different behaviors: swollen (random) coil, θ-like polymer, and equilibrium globule. **G.** Proximity frequency vs genomic distance for wild-type S15-16 embryos (blue circles), and for simulations of a self-interacting polymer in the globule regime (green curve, *U*=0.9k_B_T), a polymer at the θ-like transition (gray dashed curve, (*U*=0.65k_B_T) and of a polymer in the swollen coil regime (solid gray curve, (*U*=0k_B_T). **H.** Proximity frequency vs genomic distance, for wild-type S15-16 embryos (blue circles), Pc mutant embryos (gray triangles), and NC14 embryos (red squares). Solid and dashed lines represent a guide to the eye for the three datasets. A plateau in proximity frequency can be observed at genomic distances higher than 1 Mb.

We designed, amplified and hybridized an oligopaint-FISH library where each of the 19 Pc loci in chr3R (Fig. 1A) was encoded by a single barcode. Barcodes were sequentially imaged as described previously (Cardozo Gizzi et al. 2020) (Materials and Methods and Figs. S2A-D for more details). Next, we calculated the mean pairwise distance and ensemble proximity frequency maps for wildtype S15-S16 embryos from a large number of cells (N=22243, Fig. 1C, Materials and Methods). To validate the method, we first compared the Hi-M proximity frequency matrix for S15-S16 embryos (calculated with a proximity threshold of T=250nm), with the publicly-available Hi-C contact matrix for S16 embryos (Ogiyama et al. 2018) (Fig. S2E). Both matrices display a similar organization, notably the presence of TADs and long-range interactions. To further test the robustness of the method, we calculated the Pearson correlation between Hi-M proximity frequencies and Hi-C contact frequencies as a function of proximity threshold and genomic distance (Fig. S2F). For proximity distance thresholds equal or higher than 250 nm, Hi-M and Hi-C data displayed similar levels of correlation (p > 0.65), thus we chose a proximity threshold of 250 nm to calculate proximity frequencies. To further validate this choice, we calculated the Pearson correlation coefficient between Hi-M matrices obtained with different proximity thresholds (between 100-400 nm). In all cases, the correlation was high (p>0.9) (Fig. S2G).

Barcodes within large Pc TADs (ANT-C, BX-C and NK-C) displayed high proximity frequencies and short pairwise distances (PWD) (Fig. 1C), consistent with previous observations in *Hox* TADs (Lanzuolo et al. 2007; Sexton et al. 2012; Mateo et al. 2019; Cheutin and Cavalli 2018) (Fig. 1C, black boxes). In addition, we also observed that BX-C and ANT-C, two distant *Hox* TADs, displayed preferential proximities (Fig. 1C, green arrows), as expected from previous Hi-C and microscopy reports (Bantignies, 2011; Sexton et al. 2012). Interestingly, long-range proximities between other PcG targets were inhomogeneous, indicating that not all PcG targets have the same probability to interact with each other.

Long-range contacts between Pc targets rely on PcG proteins (Bantignies, 2011). Thus, 3D proximity should decrease in embryos lacking essential components of the PcG machinery. We tested this hypothesis by performing Hi-M in homozygous Pc mutant embryos. These embryos showed a loss of 3D proximity both at short-range (within *hox* TADs, Fig. 1D, yellow arrows) and at long-ranges (e.g. between distant Pc barcodes, Fig. 1D, green arrow), indicating that 3D proximity between Pc barcodes requires PcG proteins.

Next, we reasoned that long-range 3D proximity between Pc barcodes should be reduced before the establishment of Polycomb repression programs during development. To test this, we imaged the organization of Pc barcodes in nuclear cycle 14 (NC14/stage 5) embryos. Interestingly, barcodes within *Hox* TADs were already proximal in this early developmental stage (Fig. 1E, black boxes). However, long-range proximities between Polycomb barcodes were drastically depleted (Fig. 1E, white dashed squares), consistent with the local organization of Polycomb targets into TADs preceding the establishment of 3D long-range Polycomb contacts.

### Polycomb targets display weak polymer microphase-separation behavior

To shed light into the mechanisms responsible for the chromosome-wide organization of PcG targets observed by Hi-M, we resorted to a modeling approach that implements a lattice block copolymer model (Jost et al. 2014) (Fig. 1F, and Methods). In short, the chromosome was modeled by 866 beads with two possible identities: Polycomb or not Polycomb. The size of beads in the simulation (20kb) was slightly larger than the genomic size of Pc barcodes in our experiments (15kb). The genomic distribution of Pc beads mirrored the location of PcG targets in chromosome 3R used in Hi-M experiments (Fig. 1A), with all intervening beads labeled as non-Pc, as suggested by previous modeling studies (Jost et al. 2014). The phase diagram was established by calculating two estimators as a function of the interaction strength between Polycomb monomers (U): the internal energy *E* characterizing the average number of contacts between two Pc beads and the squared radius of gyration (*R^2^_g_*) characterizing the spatial extension of the polymer (mean square distance of the monomers with respect to the center of mass, see Methods). The polymer displays three different regimes: random coil (*U < 0.65k_B_T*), θ-like polymer (*U ~ 0.65 k_B_T*), and equilibrium globule (*U > 0.65 k_B_T*) (Figs. 1F, S2H-I) (Lieberman-Aiden et al. 2009; Mirny 2011; Halverson et al. 2014). In the random coil regime, the proximity frequency *P(s)* between monomers separated by a genomic distance s scales as *P(s) ~s*^-*3*⍰^ where ⍰ ~ 0.588 is the Flory exponent of a (non-interacting) self-avoiding polymer (Fig. 1G, gray solid curve). In the globule regime, beads attract each other and the proximity frequency between monomers scales as *P(s)* ~ *s*^-3/2^ before reaching a plateau at genomic distances corresponding to the boundary of the globule (Fig. 1G, green curve). At the so-called θ-point, the fluctuations in *E* and *R_g_* display a maximum (at the critical energy *U_c_* ~ 0.65 k_B_T for our model) (Fig. S2H-I and Methods), and the polymer represents an intermediate regime where attractive interactions between beads compensate exactly for the swelling due to self-avoidance, leading to Gaussian statistics with the proximity frequency between monomers scaling as *P(s)* ~ *s*^-3/2^ at all scales (Fig. 1G, gray dashed curve). In our particular case, the θ-like transition has to be considered with caution, due to the inhomogeneous distribution of Pc beads: high density regions of Pc beads are already in the (equilibrium) globule regime while regions of low Pc bead densities are still in the swollen coil regime (see the inhomogeneities of the proximity frequency in Fig. 1G, for the θ-like polymer).

To determine the regime that best describes the 3D folding of PcG targets in chr. 3R, we plotted the experimental proximity frequency as a function of genomic distance *P(s)*. The experimental data for S15-S16 WT embryos was best represented by an equilibrium globule (*U* = 0.9k_B_T, Fig. 1G). Below ~1Mb, the proximity frequency decreases with genomic distance as *s*^-3/2^, then reaches a plateau at larger distances (>1Mbp). The plateau in the curve is the signature of microphase separation, characteristic of block copolymers (Nuebler et al. 2018; Khokhlov, Grosberg, and Pande 1994; Rubinstein and Colby 2003; Jost et al. 2014). A block copolymer exhibits microphase separation when the two components (Pc and non-Pc beads) are non uniformly distributed in space, and display regions of local spatial enrichment where distant Pc loci may concentrate (Leibler 1980; Haddad, Jost, and Vaillant 2017). We stress, however, that the energy required to simulate the equilibrium globule that best represents the experimental data is close to the thermal energy (k_B_T), thus indicating very weak microphase separation.

We then sought to determine which polymer folding regime best described the experimental *P(s)* for NC14 embryos. Interestingly, for NC14 the *P(s)* curve was also best represented by an equilibrium globule but with a lower interaction energy (*U* = 0.8 k_B_T) (Fig. S2J). We note that this difference in interaction energy between NC14 versus S15-S16 embryos is small (0.1k_B_T) but can still lead to a large overall energy difference when integrated over the whole chromosome (containing 75 Pc beads), thus considerably impacting the global organization of the chromosome (see below). This reduction in interaction energy between S15-S16 and NC14 embryos suggests that Polycomb proteins play a role in the formation of weak microphase-separated repressive compartments.

To test this prediction, we examined the differences in experimental *P(s)* curves between S15-S16 WT, Pc mutant, and NC14 embryos. Proximity frequencies were notably reduced in the absence of Pc, and declined even rapidly in NC14 embryos, consistent with our previous conclusions regarding the role of PcG proteins in mediating long-range interactions between PcG targets (Fig. 1H). Surprisingly, *P(s)* curves for Pc depleted and NC14 embryos still exhibited a plateau above ^~^1Mb and were also well represented by an equilibrium globule (Figs. 1H, S2J-K). Furthermore, the plateau was also present when simulating the whole chromosome 3R (Figs. S2L-M), with proximity frequencies above ^~^15 Mb being extremely low (< 0.5%, Fig. S2L). Overall, these results indicate that PcG proteins reinforce the formation of weak microphase-separated compartments, but that other factors (e.g. HP1, chromatin insulators, active transcription) (Zenk et al. 2021; Hug et al. 2017; Rowley et al. 2017) are likely also involved in this process in *Drosophila*.

### Chromosome-wide association of Polycomb targets involves predominantly pairwise interactions

In *Drosophila*, Polycomb components assemble into large Pc foci (Buchenau et al. 1998; Saurin et al. 1998; Bantignies et al. 2011). These results suggest that Pc compartments may involve the spatial clustering of multiple PcG targets. To test this hypothesis, we calculated how often a Pc barcode was proximal (at a distance ≤ 250 nm) to any other Pc barcode in single cells. Targets within large Pc domains (i.e. ANT-C, BX-C, NK-C) were combined together to focus on long-range Pc contacts. In S15-S16 embryos, two or more distant Pc barcodes were found to spatially co-localize in only 4 ± 2 % of cells (Fig. 2A). This frequency was comparable for all the Pc barcodes investigated, and in all cases lower than 10%. As expected, this frequency of co-localization was even lower for NC14 embryos (2 ± 2 %) (Fig. 2B), consistent with the loss of long-range Pc proximity in early embryos (Fig. 1E), and suggesting that Pc architecture is gradually acquired during development. To determine if this behavior was dependent on the proximity distance threshold, we calculated the mean colocalization frequency of Pc barcodes for different distance thresholds (Fig. 2C). The mean proximity frequencies remained lower than 15% in most cases, even for distance cutoffs as large as 400 nm. For early embryos, the mean proximity frequencies were considerably lower (Fig. 2D). As expected, the mean co-localization frequency was positively correlated with the size of the Pc domain within which the Pc barcode was located (Fig. 2E), consistent with a role of PcG components in mediating long-range interactions between Pc domains. Thus, in single cells, Polycomb targets within large and small Pc domains rarely spatially co-localize with other Polycomb targets to form clusters.

**Figure 2:**
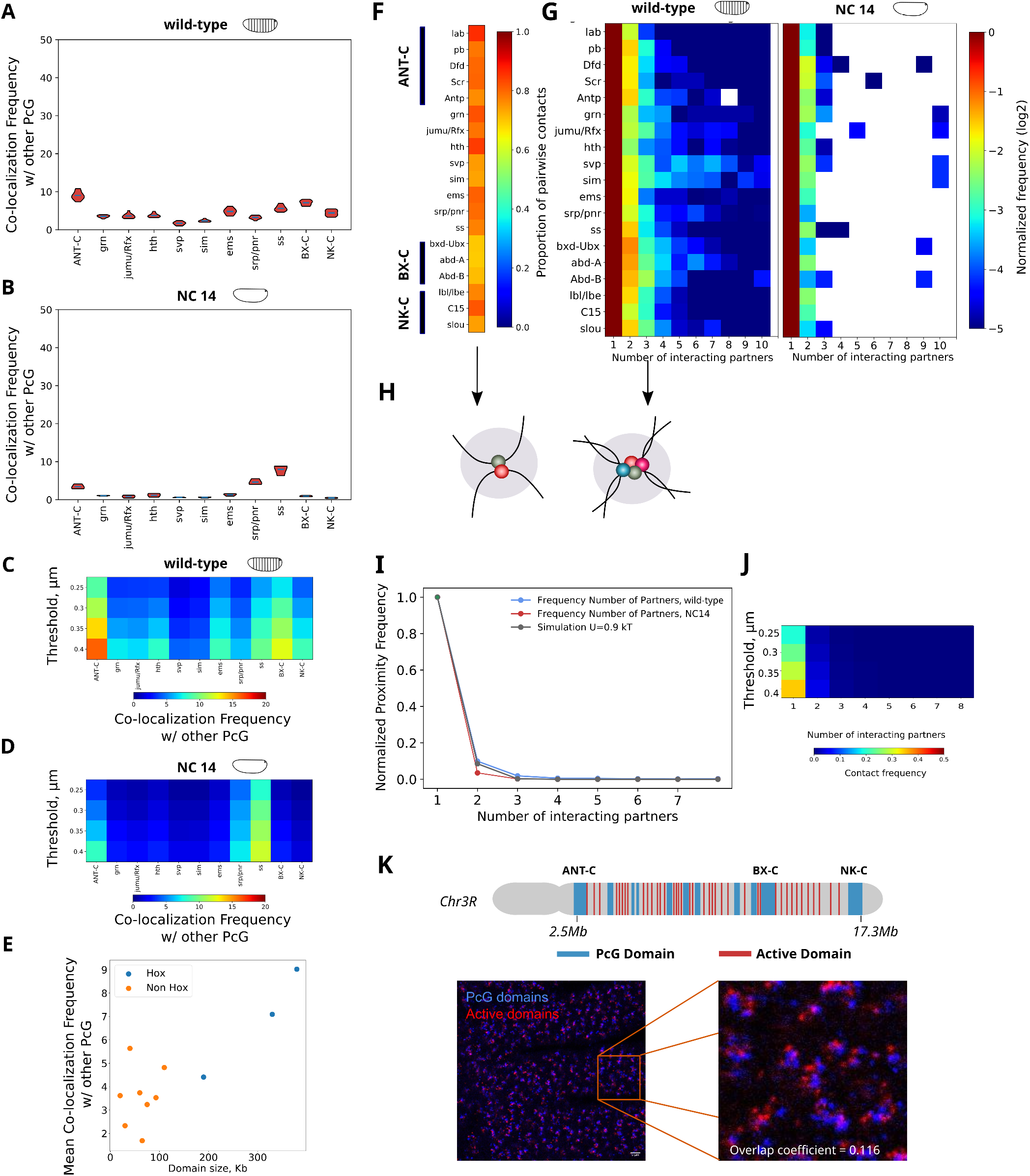
Distant Pc genes co-localize in 3D rarely, and predominantly involve only two Pc targets. **A - B.** Violin plot distributions representing the frequency with which each Pc domain interacts with any other Pc domain encoded in our oligopaint library for S15-S16 (panel A) and NC14 embryos (panel B) (see Methods). Grey line represents the mean. **C - D.** Map of mean frequency of Pc-Pc interactions (as in panels A-B) for different proximity thresholds (between 250 nm and 400 nm) for S15-S16 (panel C) and NC14 embryos (panel D). **E.** Mean inter-domain proximity frequency as a function of domain size for S15-S16 embryos. Pc domains are separated as *Hox* and *non-Hox*. Distance threshold = 250 nm. **F.** Proportion of pairwise (2 partners) versus multi-way Pc interactions (>2 partners). Pc genes are indicated on the left, colorbar represents normalized frequencies. **G.** Histograms of long-range Pc interactions as a function of the number of interacting partners, normalized by the number of pairwise interactions (1 interacting partner), for late S15-S16 (left) and NC14 (right) embryos. Pc genes are indicated on the left of the panel, and the color-scale represents normalized frequency in log2-scale. **H.** Scheme representing Pc clusters with (left panel) 1 (i.e. pairwise) and (right panel) 3 partners. **I.** Normalized proximity frequency versus number of interacting partners for wild-type S15-S16 embryos (blue curve), NC14 embryos (red curve) and simulations of a co-polymer in the equilibrium globule regime (*U =* 0.9k_B_T, gray curve). **J.** Experimental proximity frequency versus number of interacting partners for proximity thresholds between 250 nm - 400 nm (wild-type S15-16 embryos). **K.** Multiple active (red) and Polycomb domains were labeled in chr. 3R (blue) (top scheme) and then imaged together using conventional two-color confocal microscopy. Pearson’s colocalization coefficient (0.095) and overlap coefficient (0.16) were consistent with segregation of active and Pc domains. Estimation of the number of expected Pc target spots: the three *Hox* domains contain several barcodes, thus they will be detected with a high probability p = 1 - (1-p_0_)^N^ > 94%, considering a detection efficiency of p_0_ = 0.6 and a number of barcodes of N = 3. The reminder will be detected with an efficiency p_0_, thus the average number of detectable domains is ^~^ 3 + 8 * p_0_= 7.8. Two Pc domains come in close proximity, and therefore cannot be spatially resolved, at a frequency of ^~^ 0.06 (Fig. S3F). Thus we expect to detect, in average, 7.8 - 0.06 * 7.8 ^~^ 7.3 domains.

To explore whether these infrequent spatial encounters involved multiple Polycomb targets, we calculated the proportion of clusters containing two (*i.e*. pairwise cluster) or more Polycomb targets (multi-way cluster). Clusters containing only two Polycomb targets were the most common in all cases (>70%) (Fig. 2F). Next, we calculated the frequency of multi-way clusters as a function of the number of targets in a cluster, normalized by the pairwise cluster frequency (Fig. 2G). This normalized frequency of multi-way interactions decreased monotonically with the number of co-localizing targets, inconsistent with Pc foci arising from the nucleation of multiple Polycomb targets. Notably, this behavior was not tissue-specific, as the trend was similar for all segments of the embryo (Fig. S3A). For NC14 embryos, Polycomb clusters contained almost exclusively two targets, with the frequency of multi-way clusters being almost negligible (<5%, Figs. S3B, 2G). All in all, these results suggest that Polycomb targets rarely form clusters, and when they do, the cluster contains a very limited number of targets (Fig. 2H). These conclusions are in full agreement with our previous results indicating that the tendency of Pc to form microphase-separated compartments is weak.

To further test these conclusions, we calculated the frequencies with which multiple Polycomb targets co-localized in the block copolymer model presented above (*U* = 0.9 k_B_T) (Fig. 1F) and compared them to the experimental frequencies (Figs. 2I-J). The co-localization frequency between Pc beads in the model was ^~^10%, similar to our experimental measurements (Fig. 2A). In addition, the relative frequencies of pairwise and multi-way clusters were also comparable in experimental and simulated data (Fig. 2I). Thus, our simple polymer model reproduces the low experimental frequencies of pairwise and multi-way proximities, suggesting that the entropy of the polymer dominates over the enthalpic contributions provided by attractive interactions between Polycomb targets.

To further validate this hypothesis, we devised a toy polymer model containing two Polycomb targets (one in each end) and characterized its behavior for different Polycomb interaction energies (U) and polymer lengths (L) (Fig. S3D). We found that the interaction energy required to bring Polycomb beads together increased logarithmically with polymer length (Fig. S3E), while in a first approximation, the entropy barrier to bring the two Pc beads together scales, at leading order, like the log of the chromatin length. Thus, the entropy of chromatin acts to counteract the tendency of Polycomb targets to coalesce in space, providing a rationale for the weak microphase separation behavior observed for the full polymer.

Finally, we designed and imaged an oligopaint library labeling the 19 Polycomb targets in chr. 3R (Fig. 1A, shown in blue in Fig. 2K) as well as the 39 active regions between them (Fig. 2K, red; see Methods). In most cells, Polycomb and active domains were spatially segregated (Fig. 2K, overlap coefficient: 0.1, Pearson’s correlation coefficient: 0.095). From the genomic distribution of Polycomb targets in the oligopaint library, a maximum of 11 Polycomb domains should be resolvable (Fig. S3C). This estimation is lower than the total number of Polycomb targets in our design (19), as targets in close genomic proximity (i.e. inside ANT-C, BX-C and NK-C) would appear as single diffraction-limited spots. Considering our efficiency of barcode detection, and the probability of pairwise co-localization of two Pc domains, we estimate that we should be able to visualize ^~^7.3 barcode spots per cell (see legend of Fig. 2K). Remarkably, we observed 7 ± 2 Polycomb target spots per cell, consistent with a low degree of spatial clustering. All in all, these data and simulations indicate that spatial coalescence of distant Polycomb targets is limited.

### Gene repression and expression change the 3D internal organization of Polycomb domains

Our data shows that interactions between distant Polycomb targets are rare and involve primarily two targets. To determine whether these interactions depend on the transcriptional status of the co-localizing Polycomb targets, we mapped proximity frequencies for different segments of the embryo displaying distinct transcriptional programs. First, we focused on intra-domain interactions within *hox* TADs. These TADs contain the genes responsible for the development of body segments, and display well-defined patterns of expression and repression along the antero-posterior axis of the embryo (E. B. Lewis 1978; Dessain and McGinnis 1993; Edward B. Lewis et al. 2004; Kaufman, Seeger, and Olsen 1990).

We profited from the ability of Hi-M to maintain spatial information to calculate the intra-TAD proximity frequencies between *hox* target genes within BX-C and ANT-C for each segment of the embryo (Fig. 3A), and relied on existing transcriptional data to identify the segment where each gene was expressed (Fig. S4A). Proximity frequencies were normalized with respect to the segment of repression of the anchor to detect whether the expression of a Polycomb target gene changed the frequency with which it co-localized with the other *hox* genes within its TAD (Fig. 3A). For BX-C, intra-TAD normalized proximity frequencies were negative, indicating that gene repression consistently led to higher colocalization frequencies for the *hox* genes within BX-C (Figs. 3A-B). This result is consistent with previous observations (Lanzuolo et al. 2007; Cheutin and Cavalli 2018). ANT-C displayed a more complex behavior, with targets exhibiting a small negative change (*Antp*), no overall change (*lab, Scr*), or even positive changes (*pb, Dfd*) when normalized by their segment of repression (Figs. 3A and S5A). Thus, we conclude that repression of *hox* genes does not always lead to the most compact TAD configuration, perhaps due to the existence of more compact TAD conformations in segments where a subset of Pc targets are expressed (see next paragraph).

**Figure 3:**
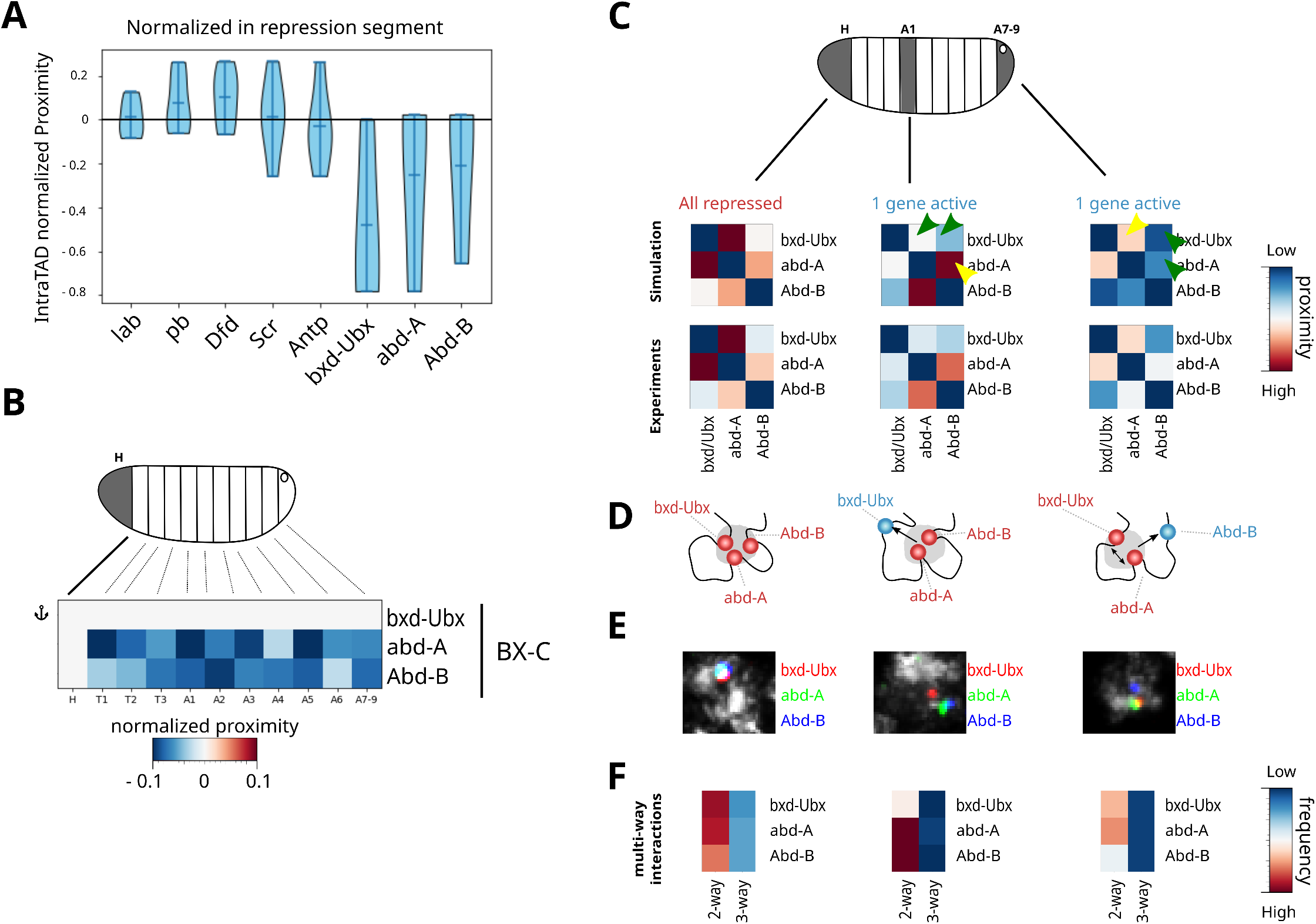
Regulation of gene expression changes chromatin organization within Hox Pc TADs. **A.** Violin plots displaying the normalized proximity frequency distributions for each Hox gene within chr. 3R with respect to the repression segment (A7-A9 for ANT-C, H for BX-C). Only intra-TAD proximities were considered. Normalized proximity frequencies were calculated by subtracting the proximity frequency of the segment minus that of the segment of repression. Violin plots were created by combining the normalized proximity frequencies for all segments for each anchor gene. **B.** Intra-TAD normalized proximity frequency maps for bxd-Ubx, normalized in the segment where all Hox genes in BX-C are repressed (head). **C.** Top: diagram showing segments where PcG genes within BX-C are all repressed (head), or where at least one target is active (A1, A7-9). Middle matrices: Simulated and experimental proximity frequency maps for Hox genes within BX-C in head, A1, and *A7-9*. Yellow arrows represent regions displaying higher proximity frequencies, and green arrows regions with lower proximity frequencies. **D.** Diagram representing the segment where proximity between Hox genes is highest (head), and segments where the gene being expressed lost proximity to the other Hox genes within the TAD (bxd-Ubx in A1 and Abd-B in A7-9). **E.** Representative microscopy images for the three Hox genes within the BX-C domain described in panel D. **F.** IntraTAD proximity frequencies for BX-C in the segments highlighted in panel D. In all segments, the frequency of pairwise interactions is predominant, and diminishes upon gene activation. The frequency of 3-way interactions is the highest for the head, where all genes are repressed.

Next, we tested whether the simple copolymer model proposed above (Fig. 1F) was able to qualitatively reproduce these observations. For this, we performed simulations under three scenarios: (1) all three genes within BX-C are repressed (head). In this case, the three genes in BX-C interact with energy *U*; (2) *bxd-Ubx* is expressed and *abd-A/Abd-B* are repressed (segment A1); (3) *Abd-B* is expressed and *bxd-Ubx/abd-A* are repressed (segments A7-9). In the last two cases, repressed genes interact with energy *U*, while the region containing the active gene is considered as non-interacting. To compare results from simulations and experiments, we plotted the proximity frequency matrices for BX-C for the head, segment A1 and segments A7-9 (Fig. 3C). Notably, the simulations were able to reproduce experimental data in the three different segments. In the head, where all genes within BX-C are repressed, proximities between Polycomb targets were high, notably between *bxd-Ubx* and *abd-A*, and *bxd-Ubx* and *Abd-B*. In segments A1/A7-9, Polycomb targets in active regions were less often proximal to repressed targets (Fig. 3C, green arrows, and Figs. 3D-E). This phenomenon can also be seen by plotting the proximity frequencies normalized by the segment of expression for each gene in BX-C (Figs. S5B). Notably, in these segments the proximity frequency between repressed genes was enhanced, likely triggered by the loss of interactions with the activated target within the TAD (Fig. 3C, yellow arrows, and Figs. 3D-E). Thus, tuning of the epigenetic state of Polycomb target genes within a TAD in the lattice copolymer model was enough to qualitatively reproduce the trends in the experimental proximity maps of cell types with different TAD configurations.

We previously established that clusters of Polycomb targets involved predominantly two genes (Fig. 2). To determine if this property depended on cell type or epigenetic state, we analyzed the distribution of multi-way interactions in the head, A1, and A7-9 segments (Fig. 3F). In all segments, the frequency of pairwise interactions was predominant, and diminished upon gene activation, consistent with our previous results. The frequency of 3-way interactions was highest for the head, where all genes are repressed. Overall, these results indicate that formation of higher-order complexes involving multiple Polycomb targets (more than two) within a TAD are modulated by epigenetic state, but remain rare, even in segments where all Polycomb targets within the TAD are repressed.

### Chromosome-wide 3D physical proximity between Polycomb domains increases in both repressed and co-expressed segments

Next, we investigated whether the co-localization of Polycomb targets located in different TADs also correlated with their transcriptional state. For this, we calculated the normalized proximity frequency between *Hox* genes and all other Polycomb targets in chr3R. Proximities were normalized to segments in which the target genes are maximally expressed (Fig. S4B). For most Polycomb targets, the normalized proximity frequency displayed positive values (Fig. 4A-C), thus *Hox* genes co-localized more often with other Polycomb targets in segments where they were repressed. Overall, these results show that activation of *Hox* genes not only leads to their local spatial segregation from other Polycomb genes within their TAD (Fig. 3), but also from more distant Polycomb targets (Figs. 4A-C, S6A-C).

**Figure 4:**
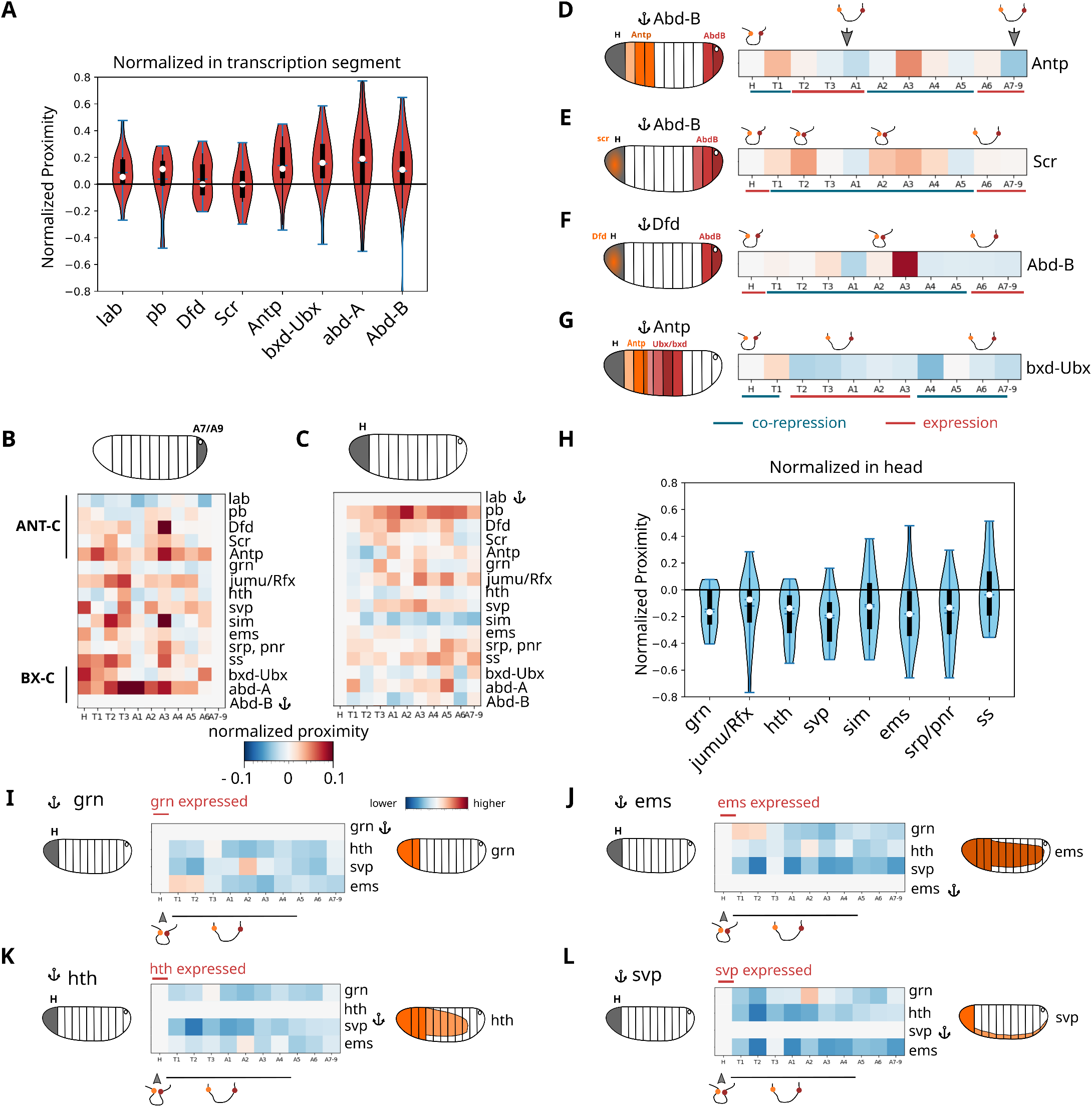
Pairwise, long-range chromatin interactions between Pc domains correlate with expression patterns. Co-expressed and co-activated genes display an increase in physical proximity. **A.** Violin plots displaying the normalized proximity frequency distributions for each Hox gene within chr. 3R. All Pc barcodes were used in the analysis, thus including intra and interTAD proximities. Normalized proximity frequencies were calculated by subtracting the proximity frequency of the segment minus that of the segment of expression. Violin plots were created by combining the normalized proximity frequencies for all segments for each anchor gene. **B - C.** Normalized proximity frequency maps for different anchors and segments. Anchors were selected at *Abd-B* (left panel) and *lab* (right panel). Proximity frequencies were normalized with respect to the segment of expression of the anchor: A7-9 (left panel) or head (right panel). **D - G.** Segment-specific normalized proximity frequency maps for long-range interactions for a selection of anchors (Abd-B, Dfd, Antp) and targets (Antp, Scr, Abd-B, bxd/Ubx). Proximity frequencies were normalized with respect to the head (gray shade in scheme). Approximate patterns of expression of anchors and targets are shown in red or orange on the schemes (left panels). Drawings above the maps represent changes in looping interactions between anchors and targets for different segments. Horizontal lines below the maps highlight the segments where the anchors and targets are co-repressed (blue), or where one of them is expressed (red). **H.** Violin plots displaying the normalized proximity frequency distributions of non-Hox target genes with all other Pc barcodes in chr. 3R. Proximity frequencies were normalized by the segment of expression of each non-Hox gene. **I - L.** Normalized proximity frequency maps for different non-Hox targets and anchors: *grn* (I), *ems* (J), *hth* (K) and *svp* (L). Proximity frequencies were normalized with respect to a segment displaying anchor expression.

Next, we explored how co-localization frequency changed with the transcriptional status of both anchor and target. For this, we analyzed the changes in proximity maps for different anchors and targets, in all cases normalized by the head, where most of the Hox genes are repressed (Figs. 4D-G). We observed that co-localization frequency between two distant Polycomb targets was highest in segments where both genes were repressed, and diminished in segments where one or the other gene was activated (Figs. 4D-G). For instance, proximity between *Abd-B* (anchor) and *Antp* (target) diminished between T2-A1, where *Antp* is active, and increased in A3-A4, where both targets are repressed (Fig. 4D). In the tail (A7-9), expression of *Abd-B* and other genes within BX-C is correlated to an overall loss of proximity between *Abd-B* and *Antp*. Similarly, *Abd-B* and *Scr/Dfd* displayed the highest proximity frequency in segments where they are both repressed (Fig. 4E-F). A similar behavior was observed for *Antp* (anchor) and *bxd-Ubx* (target) (Fig. 4G). Thus, spatial colocalization between distant *Hox* Polycomb targets was highest in segments where both targets are repressed, and was lower in segments where one of the targets was activated.

Finally, we tested whether *non-Hox* Polycomb target genes displayed a similar behavior. For this, we calculated the proximity maps for the non-Hox targets displaying clear antero-posterior expression patterns: *ems, hth, svp* and *grn* (Figs. 4H and S4B). We normalized proximity maps by the proximity frequencies of the segment where the anchor exhibited maximal expression (Fig. 4H). Notably, normalized proximity frequencies were negative for the majority of segments and targets (Fig. 4H and 4I–4L). Thus, and in contrast to *Hox* polycomb targets, non-Hox Polycomb targets displayed the highest proximity frequencies in regions where they are co-expressed. All in all, our results show that proximity frequencies of distant Polycomb targets are spatially modulated, and can be tuned in different cell types by co-repression as well as by co-expression.

## Discussion

Previous evidence suggested that multiple Pc TADs may often associate with each other in single cells to form Pc compartments (bodies) to reinforce gene repression (Isono et al. 2013; Delest, Sexton, and Cavalli 2012; Bantignies and Cavalli 2011). Here, we investigated the nature of these compartments by implementing a multiplexed imaging-based approach that maps the multiscale organization of Polycomb target genes in different presumptive tissues within the Drosophila embryo.

Pairs of Polycomb target genes are able to interact in 3D by ‘gene kissing’, an activity that requires Polycomb components and possibly other factors, such as chromatin insulators (Bantignies et al. 2011; Sexton et al. 2012). The ability of our approach to detect multiple Polycomb target genes in single cells allowed us to shed light into the nature of these kissing interactions. Previous studies determined that interactions between pairs of distant *Hox* genes were rare (10-20%), and proposed that the existence of multiple accessible Polycomb partners may explain why two Hox genes only interact in a small fraction of nuclei (Bantignies et al. 2011; Sexton et al. 2012). In fact, our analysis shows that spatial colocalization of any two distant intra-chromosomal Polycomb targets is rather infrequent. These results are consistent with a recent single-nucleus Hi-C study in *Drosophila* BG3 cells showing a weak enrichment in long-range interactions between Pc repressed regions (Ulianov et al. 2021). Moreover, we show that these rare long-range interactions are acquired after the emergence of TADs at NC14 (Hug et al. 2017; Ogiyama et al. 2018), and concomitantly with the enrichment of Pc foci (Cheutin and Cavalli 2012). Notably, our data shows that frequencies of long-range interactions vary widely between targets and do not only depend on genomic distance, suggesting a role for other factors (e.g. insulators) in the modulation of interaction specificity.

Previous genome-wide studies showed extensive interactions between distant Polycomb genes (Bantignies et al. 2011), raising the possibility that Polycomb repressive compartments could involve the coalescence of multiple repressed genomic regions. We tested this hypothesis by directly calculating the frequency of pairwise versus multi-way interactions. Notably, we found that binary interactions are predominant, with the frequency of multi-way contacts drastically decreasing with the number of targets. This finding is consistent with previous imaging studies showing that Polycomb TADs often appear as discrete 3D chromosomal units (Szabo et al. 2018) and with recent single-nucleus Hi-C work indicating that Polycomb TADs interact with each other in a cell-specific manner (Ulianov et al. 2021). Overall, these results indicate that Polycomb repressive compartments most often contain only two Polycomb target genes.

This result is supported by polymer modeling, which shows that a weak microphase-separated polymer in the globule regime correctly captures the behavior of Polycomb domains. This polymer is close to the θ-like transition in the phase diagram. In this configuration, a small change in the interaction energy between monomers leads to a large change in the overall energy of the polymer, allowing chromatin to switch conformation easily with a small difference in interaction energies. Weak microphase separation is consistent with the dynamic occupancy of Pc sites by PRC1 proteins, observed both in *Drosophila* and mammals (Ficz, Heintzmann, and Arndt-Jovin 2005; Fonseca et al. 2012; Steffen et al. 2013; Huseyin and Klose 2021), and with a recent singlenucleus Hi-C study showing that long-range Pc contacts occur regardless of their genomic distance (Ulianov et al. 2021). Furthermore, this behavior is maintained in *Pc* mutants, consistent with other factors (HP1, insulators, transcription hubs, or other PcG proteins subunits) also likely contributing to the long-range organization of Pc domains (Zenk et al. 2021; Hug et al. 2017; Rowley et al. 2017; Ulianov et al. 2021; Tuszynska, Bednarz, and Wilczynski 2021), and providing a scaffold that facilitates the encounter of distant Pc domains. This polymer model correctly captures the behavior of Pc domains, notably the predominantly pairwise nature of interacting partners, and the correlation between Polycomb architecture and gene expression. Overall, our experiments and simulations suggest that Polycomb repressive compartments form by infrequent associations of Polycomb domains. PRC1 proteins play an important role in the formation of these compartments, however, other factors such as the entropy of the chromatin polymer, specific contacts mediated by other chromatin factors, and attractive interactions between active or repressed regions are also relevant. This suggests that the composition of Polycomb compartments could be regulated by their epigenetic and transcriptional status.

We tested this hypothesis by resorting to the ability of our method to reconstruct chromatin architecture in embryonic segments with different epigenetic and transcriptional states. Remarkably, we found that proximity frequencies between Polycomb targets are modulated by both transcriptional repression and activation. *Hox* genes were colocalized most frequently in segments where they were co-repressed, both for targets located within the same TAD or for very distant genes. This result was consistent with previous observations on a limited number of targets (Lanzuolo et al. 2007; Bantignies et al. 2011; Cheutin and Cavalli 2018). Notably, transcriptional activation of *Hox* genes led to their spatial segregation, locally from other Polycomb targets within their TAD, and more globally from other repressed distant Polycomb targets. Finally, *non-Hox* genes more frequently colocalized in regions where they were both expressed, consistent with previous observations on a limited number of targets (Li et al. 2013). These interactions among coexpressed genes might depend on trithorax-group factors that can physically interact with Polycomb components to activate gene expression (Strübbe et al. 2011; Kadoch et al. 2017; Stanton et al. 2017).

In conclusion, our data are inconsistent with repressive Polycomb compartments being formed by the extensive coalescence of multiple distant Polycomb regions, and instead show that interactions between Polycomb genes occur infrequently, and involve mostly pairwise encounters modulated by transcriptional status.

## Acknowledgements

This project was funded by the European Union’s Horizon 2020 Research and Innovation Program (Grant ID 724429) (M.N.). We acknowledge the Bettencourt-Schueller Foundation for their prize ‘Coup d’élan pour la recherche Française’, the France-BioImaging infrastructure supported by the French National Research Agency (grant ID ANR-10-INBS-04, “Investments for the Future”), and the Drosophila facility (BioCampus Montpellier, CNRS, INSERM, Univ Montpellier, Montpellier, France).

## Author Contributions

Conceptualization, J.G., F.B. and M.N; Methodology, J.G., C.H., J-B.F, J-C.W., and M.N.; Investigation, J.G., J-C.W., F.B., and M.N.; Writing – Original Draft, J.G. and M.N.; Writing – Review & Editing, J.G., J-C.W., J-B.F., M.S., G.C., F.B., and M.N.; Funding Acquisition, M.N.; Resources, J.G., C.H., J-B.F., M.S., F.B.; Supervision, F.B. and M.N.

## Methods

### Probe selection and library design

A portion of ^~^15 Mb of chromosome 3R was selected. A 3 node self-organizing map (SOM, ‘kohonen’ R package) was used to produce a 3-way segmentation of 10 Kb genome wide bins. Each bin was scored based on the average ChIP-seq read counts of H3K27me3, H3K4me3 and H3K36me3 from 14-16 hr embryos (modEncode, 3955 H3K27me3, Embryos-14-16 hr, OregonR, ChIP-seq; modEncode, 5096: H3K4me3, Oregon-R, Embryos 14-16 hr OR, ChIP-seq; 4950: H3K36me3, Oregon-R, Embryos 14-16 hr OR, ChIP-seq). Each SOM node was treated as a discrete cluster and contiguous bins assigned to the same node were merged into one epi-domain. Only epi-domains of a size equal or bigger to 20 kb were selected. For Pc domains, these epi-domains were later reselected based on the enrichment of H3K27me3 (modEncode, 3955 H3K27me3, Embryos-14-16 hr; OregonR;ChIP-seq), and of PRC1 subunits PC, PH for embryos of 16 - 18 hrs of development (coming from Schuettengruber B, et. al, 2014, accession number GSE60428) and PSC (modEncode, 3960: Psc;Oregon-R;Embryos 14-16 hr OR;ChIP-seq, D. melanogaster). Domains having peaks of at least two of these PRC1 subunits were kept. They were also visually inspected using Hi-C maps from (Ogiyama et al. 2018). For active domains, they were re-selected based on the enrichment of H3K4me3 and H3K36me3 (modEncode, 5096: H3K4me3;Oregon-R;Embryos 14-16 hr OR;ChIP-seq; 4950: H3K36me3; Oregon-R; Embryos 14-16 hr OR; ChIP-seq), and they were also visually inspected using Hi-C maps from (Ogiyama et al. 2018). For inactive domains, they were selected based on the absence of the aforementioned epigenetic marks. Domains between 20 - 100 Kb were labeled by one 15 Kb probe, centered in the middle of the domain, for all domain types. Domains between 100 - 200 Kb were labeled by two 15 kb probes centered at PcG protein peaks for Pc domains, and two 15 Kb domains homogeneously distributed for active and inactive domains. Domains bigger than 200 Kb comprise ANTP-C, BX-C and NK-C, and are labeled by 15 Kb probes targeting the promoters of their genes (5,3 and 3 probes respectively). The selected Polycomb targets correspond to the PcG targets in Chromosome 3R displaying the large majority of interactions (Bantignies, 2011). Polycomb domains located outside the selected ^~^15 Mb region, or smaller than 20 Kb were not labeled as they interacted with other Polycomb genes very infrequently.

### Library synthesis and amplification

The library synthesis method is based on the development by (Beliveau et al. 2012; 2015). After selecting the genomic regions of interest, a database of genomically unique, non overlapping sequences was used to generate the Oligopaint primary probes (Oligopaints website, https://oligopaints.hms.harvard.edu/). They were mined using OligoArray (Rouillard, Zuker, and Gulari 2003). Each Oligo of the library is made of 148 nucleotides (nt), and consists in (from 5’ to 3’): a 22-nt forward universal primer region for library amplification, a 20-nt readout region, unique for each 15 kb targeted region, a 2-nt spacer, a 20-nt readout region, unique for each 15 kb targeted region, a 42-nt region of homology to chromosomal DNA, a 20-nt readout region, unique for each 15 kb targeted region, and a 22-nt unique reverse primer for library amplification. An Oligopool with all the nucleotides used was ordered from Custom Array.

The procedure to amplify the library consists in four main steps: I) PCR amplification of the Oligopaints library using a reverse primer that adds the T7 promoter sequence; II) Conversion of the PCR product to RNA via an in vitro transcription using T7 polymerase; III) Generation of singlestranded DNA (ssDNA) via reverse transcription; IV) Degradation of the RNA template using alkaline hydrolysis. The full protocol can be found at (Cardozo Gizzi et al. 2020).

Barcode sequences can be found in Supplementary Tables 1-2.

Primer sequences for library amplification are listed below (5’ to 3’):

BB291.fw: CAGGTCGAGCCCTGTAGTACG

BB292.rev: GTGTCCGAGGCTGTCTCCTAG

### Adaptor, displacement and imaging oligos

Adaptor oligos were used to bind the fluorescent label to the primary Oligopaint library. They’re made of 62 base pairs, and consist in (from 5’ to 3’): I) a binding region of 20 bp, complementary to the binding site of the oligos in the primary library; a bridge of 10 bp; and a binding site for the imaging oligo, of 32 bp.

The bridge is used to bind a displacement oligo. A displacement oligo consists in the complementary sequence of the bridge, followed by the complementary sequence of the binding region on the adaptor oligos. When injected at the proper concentration, they bind the adaptor oligo. The adaptor oligo bound by a displacement oligo can then be washed out, removing with it the fluorophore in the imaging oligo. This technique was first used in Mateo et. al. 2019, and it is used in our work to remove the fluorescent signal from the fiducial mark every 10 cycles, to avoid bleaching and to ensure an optimal signal intensity. To remove the fluorescent signal from barcodes, a chemical bleaching step was performed (see image acquisition section).

Imaging oligos consist of a 32-mer complementary to the binding region in the adaptor oligo, followed by a cleavable Alexa-647 fluorophore. Fiducial marker imaging oligo consists of a 32-mer complementary to the binding region of the corresponding adaptor oligo, followed by a non-cleavable Rhodamine Red fluorophore. Adaptors, displacement and imaging oligos were synthesized by Integrated DNA Technologies (IDT; Coralville, USA). Their sequences can be found in Supplementary Table 3.

### Embryo collection and fixation

Oregon-R w^1118^ flies were used for the WT strain. For the mutant line, the *Pc* strain was used. It consists in a null mutant (Franke, Messmer, and Paro 1995) that was balanced over the KrGFP-TM3 Sb balancer (TKG: obtained from BL#5195 of the Bloomington Drosophila Stock Center). Flies were maintained at room temperature with natural light/dark cycles and were grown on standard cornmeal yeast media at 21°C.

Following a pre-laying period of 16-18 H in cages with yeasted 0.4% acetic acid agar plates, agar plates were changed for new ones so flies can lay eggs during the corresponding time (1.5 H for NC14 embryos, 4 H for S15-16 embryos and mutants) on the new plates. Embryos were then incubated at 25°C for the corresponding time to obtain the desired developmental stage for fixation (1 H for NC14, 12 H for S15-16 and mutants). For fixation, embryos were dechorionated with bleach for 5 min and thoroughly rinsed with water. They were fixed in a fixation buffer (1:1 mixture of 4% methanol-free formaldehyde in PBS and heptane) by agitating vigorously for 15 s and then letting stand the vial for 25 min at RT. The bottom formaldehyde layer was replaced by 5 mL methanol and embryos were vortexed for 30 s. Embryos that sank to the bottom of the tube, devitellinized, were rinsed three times with methanol. Embryos were stored in methanol at −20°C until further use.

### Hybridization of Hi-M library

Embryos were rehydrated by incubating them sequentially for 5 min at RT on a rotating wheel, using 1 mL of each of the following solutions: I) 90% methanol, 10% PBT; II) 70% methanol, 30% PBT; III) 50% methanol, 50% PBT; IV) 30% methanol, 70% PBT; V) 100% PBT. Then embryos were RNAse treated during 2h at RT, and permeabilized in 0,5% Triton/PBS during 1h. Next, embryos are incubated for 20 min at RT on a rotating wheel sequentially in the following Triton/pHM solutions (pHM: 2X SSC, NaH2PO4 0.1M pH = 7, 0.1% Tween-20, 50% formamide (v/v)): I) 80% PBS-Triton, 20% pHM; II) 50% PBS-Triton, 50% pHM; III) 20% PBS-Triton, 80% pHM; IV) 100% pHM. Then 225 pmol of primary probe were diluted in 25 μL of FHB (FHB =50% Formamide, 10% dextran sulfate, 2X SSC, Salmon Sperm DNA 0.5 mg mL). Primary probes and embryos were denatured by incubating them 15 min at 80 °C. Embryos were then transferred to a 500 μL PCR tube, next pHM was removed from embryos and 30 μL of the denatured probes were added. Embryos were then placed into a thermocycler with the following program: Starting from 80°C, 43 cycles of 10 minutes, with a temperature drop of −1°C/cycle, then incubation at 37°C indefinitely. Embryos were then transferred back to a 1.5 mL tube, and sequentially washed for 20 min in the following solutions: I) 50% (vol/vol) formamide, 2× SSC; repeat this wash once; II) 40% (vol/vol) formamide, 2× SSC; III) 30% formamide, 70% PBT; IV) 20% formamide, 80% PBT; V) 10% formamide, 90% PBT; VI) 100% PBT; VII) 100% PBS-Triton. Washes I-IV were performed at 37 °C in a thermal mixer with agitation (900 r.p.m.); washes V–VII were performed at RT on a rotating wheel. An additional crosslink in 4% PFA was performed. Embryos were washed and resuspended in PBS, and stored at 4°C until use.

### Microscope setup

Experiments with NC14 embryos were performed on a home-made wide-field epifluorescence microscope built on a RAMM modular microscope system (Applied Scientific Instrumentation) coupled to a microfluidic device as described previously (Cardozo Gizzi et al. 2019; 2020). Samples were imaged using a 60x Plan-Achromat water-immersion objective (NA = 1.2, Nikon, Japan). The objective lens was mounted on a closed-loop piezoelectric stage (Nano-F100, Mad City Labs Inc. - USA). Illumination was provided by 3 lasers (OBIS-405/640 nm and Sapphire-LP-561 nm, Coherent - USA). Images were acquired using a sCMOS camera (ORCA Flash 4.0V3, Hamamatsu – Japan), with a final pixel size calibrated to 106 nm. A custom-built autofocus system was used to correct for axial drift in real-time and maintain the sample in focus as previously described (Cardozo Gizzi et al. 2019). A fluidic system was used for automated sequential hybridizations, by computer-controlling a combination of three eight-way valves (HVXM 8-5, Hamilton) and a negative pressure pump (MFCS-EZ, Fluigent) to deliver buffers and secondary readout probes onto a FCS2 flow chamber (Bioptechs). Software-controlled microscope components, including camera, stages, lasers, pump, and valves were run using a custom-made software package developed in LabView 2015 (National Instrument) and available <u>here</u>.

Experiments with S15-S16 and Pc del embryos were performed on an AxioObserver microscope coupled to a LSM800 laser-scanning confocal detection (Zeiss, Germany). Samples were imaged using a 63x, NA = 1.2 water-immersion objective (W DICII, Zeiss). Illumination was provided by 3 laser lines (405/561/640 nm). Images were acquired with a pixel size of 100 nm, and 0.5 μm z-slices. A pinhole size of 62 μm was used. ZEN 2.3/6 blue edition (.NET Framework Version: 4.0.30319.42000) was used for acquisition. Focus reproducibility during the experiment was ensured by the built-in autofocus tools available in ZEN.

Sequential hybridizations were performed using a computer-controlled fluidic system. In brief, a peristaltic pump (Instech, P720) coupled to an eight-way valve (HVXM 8-5, Hamilton) delivers the buffers into a FCS2 flow chamber (Bioptechs). Barcodes were injected sequentially using a homemade delivery platform composed of a rotating tray where the tubes are arranged (Physik Instrumente, M-404.4PD). A needle coupled to a linear stage (Physik Instrumente, VT-80) is used to inject the barcodes into the chamber. A second peristaltic pump (Instech, P720) is coupled to the needle and a two-way valve (HVXM 2-5, Hamilton), to wash the residual barcode solution from the needle between cycles. Flow rate is constantly monitored (FRP, flow-rate platform, Fluigent) in order to control the injected volumes and ensure reproducible hybridization conditions for all probes.

Finally, a XY translation stage (MS2000, Applied Scientific Instrumentation) is used to select the positions of the embryos. Pumps, valves, and translation stages were controlled using a custom-made software package developed in LabView 2015 (National Instruments). Synchronization between injections and confocal acquisitions was ensured using a trigger box (SVB-1 Zeiss, Germany) and an analog voltage output device (USB-3104, Measurement computing).

### Image acquisition

Embryos labeled with the primary library were attached to a poly-L-lysine coated coverslip, and placed into the FCS2 fluidics chamber. Fiducial mark labeling with a Rhodamine labeled readout probe and DAPI staining were performed in the chamber, using the fluidics system to inject solutions and perform washes. For image acquisition, the fluidics system harbored: 1 tube with 50 mL of washing buffer (WB, 2X SSC, 40% v/v formamide), 1 tube with 50 mL of 2x SSC, 1 tube with 20 mL of imaging buffer (IB, 1xPBS, 5% w/v glucose, 0.5 mg/mL glucose oxidase and0.05 mg/mL catalase), 1 tube with 50 mL of chemical bleaching buffer (CB, 2X SCC, 50 mM TCEP hydrochloride), and 19 tubes with 2 mL of each readout probe solution (25 nM readout probe, 2X SSC, 40% v/v formamide). To avoid degradation by oxygen, IB was stored under a layer of mineral oil throughout the experiment, and renewed every 12 h.

Several 200μm X 200μm fields of view (FOV) containing embryos were selected, using our homemade LabView software package. Z stacks of 15 - 20 μm were selected, with steps of 250 nm in the widefield setup and 500 nm in the Airyscan.

DAPI was imaged first, together with fiducial marks, using the 405, and the 561 nm laser lines. Barcode imaging was also performed automatically using our home-made Labview software package, which controlled the fluidics system and the XY translations stage and synchronized image acquisition performed by ZEN. Briefly, the chamber was filled with 1.7 mL barcode probe solution over ^~^17 min to ensure binding of readout probes. Next, the sample was washed with 1.5 mL of wash buffer for 10 min. Then 1.5 mL of 2X SSC were flushed during 10 min and finally 1.2 mL of imaging buffer was injected in ^~^12 min. Flow was stopped, and the FOVs were imaged in two channels by exciting at 561 and 641 nm to image fiducial marks and barcode probes, respectively. After imaging, the fluorescence signal of the barcode probes was cleaved using chemical bleaching by flowing 1.5 mL of CB buffer for 10 min. The Rhodamine-labeled fiducial barcode was insensitive to chemical removal. After chemical bleaching, the chamber was flushed with 1.5 mL of 2X SSC for 10 min and a new hybridization cycle started. All buffers were freshly prepared and filtered for each experiment. Barcodes displayed a labeling efficiency in the 40-65% range (Fig. S2C).

### Polymer modelling

A block copolymer of N=866 beads, each of size *a*=20kb, matching the genomic size and distribution of the experimental probes was implemented and simulated by Monte Carlo simulations (Walter and Barkema 2015) on a Face-Centered Cubic lattice (FCC). This lattice polymer model has proven to be extremely precise up to second order corrections when compared to analytical results for DNA hybridization/melting (Sakaue et al. 2017) and for the unwinding dynamics (Walter et al. 2013). Beads were divided into two classes in all simulations: Pc beads, that displayed a finite interaction strength *U*; and non-Pc beads, for which *U*=0. To study the behavior of a block copolymer, we used two estimators. The first estimator is the internal energy *E* defined as:

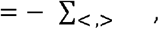

where *U* is the interaction energy between two Pc beads, is the occupation variable of the vertex of the lattice by a Pc bead (1 if a Pc bead is present and 0 otherwise) and the sum ∑_<, >_ runs over all the pairs <, > of nearest-neighbor Pc beads on the lattice. By construction, the internal energy estimates the averaged number of Pc-Pc interactions, which increases from the coil to the globule regime as *U* increases. The second estimator is the squared radius of gyration defined as:

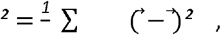

where 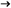 is the position vector of the monomer *i* and 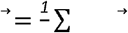 is the center of mass of the polymer. R^2^_g_ characterizes the spatial extension of the polymer, which displays a large drop from the coil to the globule regime as *U* increases. This change of the spatial conformation is the cause of the different contact probability described in Fig. 1G.

In order to assess the behavior of this polymer versus *U*, we built the phase diagram in Figs. S2H-I (main graph) where the internal energy *E* and the squared radius of gyration R^2^_g_ are plotted versus U. To do so, R^2^_g_ was sampled from U=0 over 2.10^5^ configurations. N^2^ Monte Carlo steps were performed to decorrelate the polymer between two samplings. Subsequently to the completion of the sampling at a given *U*, the value of *U* was increased by 0.05k_B_T, the system was thermalized during 10^7^ Monte Carlo steps and the sampling procedure was resumed at this new interaction energy. In the inset, we plotted the fluctuations of both *E* and R^2^_g_ that were calculated in the same manner. Different values of *U* were used, to match experimental data: for all observables presented in the main text (Frequency versus genomic distance; histogram of pairwise interactions and proximity map between ubx, abd-A and Abd-B) we used *U* = 0.8 k_B_T for early embryos and *U* = 0.9 k_B_T for late embryos.

### Data analysis

Airyscan images (.czi format) were converted to TIFF files using the Bio-Formats plugin in Fiji (Schneider, Rasband, and Eliceiri 2012; Schindelin et al. 2012, https://github.com/ome/bioformats, (Linkert et al. 2010). Raw images were deconvolved using Huygens Professional version 20.04 (Scientific Volume Imaging, the Netherlands,https://svi.nl/), via the CMLE algorithm (SNR:20, 40 iterations) with a custom-made script written in Tcl/Tk.

The following analysis steps were performed with home-made scripts written in MATLAB 2019b (The MathWorks, Inc., Natick, United States) and available in <u>Mendeley Data</u>. First, X-Y drift is corrected for each hybridization cycle. A global X-Y correction for each cycle is obtained by cross-correlating the images of the fiducial of each cycle to that of the reference fiducial (reference cycle). This produces a single 3D vector for each barcode cycle that represents a *global* correction applied to the whole FOV. Second, an adaptive thresholding is used to pre-segment the spots of each fiducial for all FOVs and for all barcode cycles. The 3D coordinates of each fiducial spot in each FOV and cycle were then estimated by segmentation and 3D Gaussian fitting. Fiducial spots with sizes larger than the diffraction limit (~2.2 pixels in our microscope) were filtered out. Third, we obtained ‘local’ 3D correction vectors for each cell in each FOV by first using the global X-Y correction vector to pre-align fiducial spots in each cycle to fiducial spots in the reference cycle. Then, image-based cross-correlation of these pre-aligned fiducial spots were used to reach subpixel accuracy in the correction vector. This approach allowed for 3D, subpixel accuracy drift-correction across the whole FOV (Fig. S2D). Fourth, barcode spots were segmented for all hybridization cycles in batch processing mode using optimized adaptive thresholding. The 3D coordinates of each barcode were then determined by 3D Gaussian fitting of the segmented regions. These positions were corrected for drift by using the closest fiducial barcode vector obtained from the previous analysis step. Nuclei were segmented from DAPI images by adaptive local thresholding and watershed filtering. Embryo’s segments were selected by manually drawing polygons over projected DAPI images. This procedure was used to assign each DAPI-segmented cell to the corresponding segment. Then, barcodes were attributed to each cell by using DAPI segmentation masks and the 3D localizations of barcodes. For each nuclei, we then calculated pairwise distance matrices. All further analysis was performed using home-made Python 3 scripts. In Figs. 1C-E, the proximity frequency was obtained as the number of nuclei where the pairwise distances were lower or equal to 250 nm, normalized by the number of nuclei containing both barcodes. Outliers (defined using the interquartile rule) or bins with no reported interactions were filtered out. In Figs. 2A-B, for each Pc anchor we calculated the frequency of cells co-localizing with at least one Pc target. This calculation was repeated for each Pc anchor in each segment of the embryo. Violin plots for each Pc target were obtained by merging together frequencies from different segments. A similar procedure was used for NC14 embryos, but in this case the segments were defined by the anterior, median, and posterior regions (with roughly similar number of cells). Image processing was carried out on Linux terminals connected to a server running Linux PopOS 19.10, with four GeForce GTX 1080Ti GPU cards (SCAN computers, UK).

## Data and code availability

Code used in this manuscript is publicly available from <u>Mendeley Data</u> (Hi-M data analysis), from our <u>HiMacquisitionSoft</u> Github repository (Labview acquisition software) and from a <u>Github Repository</u> containing scripts specific to this study. Datasets are publicly available from <u>Zenodo</u> (DOI: 10.5281/zenodo.6461558).

## References

Andrey, Guillaume, and Stefan Mundlos. 2017. “The Three-Dimensional Genome: Regulating Gene Expression during Pluripotency and Development.” Development 144 (20): 3646–58. https://doi.org/10.1242/dev.148304.

Bantignies, Frédéric, and Giacomo Cavalli. 2011. “Polycomb Group Proteins: Repression in 3D.” Trends in Genetics 27 (11): 454–64. https://doi.org/10.1016/j.tig.2011.06.008.

Bantignies, Frédéric, Virginie Roure, Itys Comet, Benjamin Leblanc, Bernd Schuettengruber, Jérôme Bonnet, Vanessa Tixier, André Mas, and Giacomo Cavalli. 2011. “Polycomb-Dependent Regulatory Contacts between Distant Hox Loci in Drosophila.” Cell 144 (2): 214–26. https://doi.org/10.1016/j.cell.2010.12.026.

Beliveau, Brian J., Alistair N. Boettiger, Maier S. Avendaño, Ralf Jungmann, Ruth B. McCole, Eric F. Joyce, Caroline Kim-Kiselak, et al. 2015. “Single-Molecule Super-Resolution Imaging of Chromosomes and in Situ Haplotype Visualization Using Oligopaint FISH Probes.” Nature Communications 6 (1): 7147. https://doi.org/10.1038/ncomms8147.

Beliveau, Brian J., Eric F. Joyce, Nicholas Apostolopoulos, Feyza Yilmaz, Chamith Y. Fonseka, Ruth B. McCole, Yiming Chang, et al. 2012. “Versatile Design and Synthesis Platform for Visualizing Genomes with Oligopaint FISH Probes.” Proceedings of the National Academy of Sciences 109 (52): 21301–6.

Bintu, Bogdan, Leslie J. Mateo, Jun-Han Su, Nicholas A. Sinnott-Armstrong, Mirae Parker, Seon Kinrot, Kei Yamaya, Alistair N. Boettiger, and Xiaowei Zhuang. 2018. “Super-Resolution Chromatin Tracing Reveals Domains and Cooperative Interactions in Single Cells.” Science 362 (6413). https://doi.org/10.1126/science.aau1783.

Boettiger, Alistair N., Bogdan Bintu, Jeffrey R. Moffitt, Siyuan Wang, Brian J. Beliveau, Geoffrey Fudenberg, Maxim Imakaev, Leonid A. Mirny, Chao-ting Wu, and Xiaowei Zhuang. 2016. “Super-Resolution Imaging Reveals Distinct Chromatin Folding for Different Epigenetic States.” Nature 529 (7586): 418–22. https://doi.org/10.1038/nature16496.

Buchenau, Peter, Jacob Hodgson, Helen Strutt, and Donna J. Arndt-Jovin. 1998. “The Distribution of Polycomb-Group Proteins During Cell Division and Development in Drosophila Embryos: Impact on Models for Silencing.” Journal of Cell Biology 141 (2): 469–81. https://doi.org/10.1083/jcb.141.2.469.

Cardozo Gizzi, Andrés M., Diego I. Cattoni, Jean-Bernard Fiche, Sergio M. Espinola, Julian Gurgo, Olivier Messina, Christophe Houbron, et al. 2019. “Microscopy-Based Chromosome Conformation Capture Enables Simultaneous Visualization of Genome Organization and Transcription in Intact Organisms.” Molecular Cell 74 (1): 212–222.e5. https://doi.org/10.1016/j.molcel.2019.01.011.

Cardozo Gizzi, Andrés M., Sergio M. Espinola, Julian Gurgo, Christophe Houbron, Jean-Bernard Fiche, Diego I. Cattoni, and Marcelo Nollmann. 2020. “Direct and Simultaneous Observation of Transcription and Chromosome Architecture in Single Cells with Hi-M.” Nature Protocols 15 (3): 840–76. https://doi.org/10.1038/s41596-019-0269-9.

Cattoni, Diego I., Andrés M. Cardozo Gizzi, Mariya Georgieva, Marco Di Stefano, Alessandro Valeri, Delphine Chamousset, Christophe Houbron, et al. 2017. “Single-Cell Absolute Contact Probability Detection Reveals Chromosomes Are Organized by Multiple Low-Frequency yet Specific Interactions.” Nature Communications 8 (1): 1753. https://doi.org/10.1038/s41467-017-01962-x.

Cavalli, Giacomo, and Tom Misteli. 2013. “Functional Implications of Genome Topology.” Nature Structural & Molecular Biology 20 (3): 290–99. https://doi.org/10.1038/nsmb.2474.

Cheutin, Thierry, and Giacomo Cavalli. 2012. “Progressive Polycomb Assembly on H3K27me3 Compartments Generates Polycomb Bodies with Developmentally Regulated Motion.” PLOS Genetics 8 (1): e1002465. https://doi.org/10.1371/journal.pgen.1002465.

Cheutin, Thierry, and Giacomo Cavalli. 2018. “Loss of PRC1 Induces Higher-Order Opening of Hox Loci Independently of Transcription during Drosophila Embryogenesis.” Nature Communications 9 (1): 3898. https://doi.org/10.1038/s41467-018-05945-4.

Delest, Anna, Tom Sexton, and Giacomo Cavalli. 2012. “Polycomb: A Paradigm for Genome Organization from One to Three Dimensions.” Current Opinion in Cell Biology, Nucleus and gene expression, 24 (3): 405–14. https://doi.org/10.1016/j.ceb.2012.01.008.

Dessain, Scott, and William McGinnis. 1993. “Drosophila Homeobox Genes.” In Advances in Developmental Biochemistry, edited by Paul M. Wassarman, 2:1–55. Advances in Developmental Biochemistry. Academic Press. https://doi.org/10.1016/S1064-2722(08)60035-3.

Dixon, Jesse R., Siddarth Selvaraj, Feng Yue, Audrey Kim, Yan Li, Yin Shen, Ming Hu, Jun S. Liu, and Bing Ren. 2012. “Topological Domains in Mammalian Genomes Identified by Analysis of Chromatin Interactions.” Nature 485 (7398): 376–80. https://doi.org/10.1038/nature11082.

Ficz, Gabriella, Rainer Heintzmann, and Donna J. Arndt-Jovin. 2005. “Polycomb Group Protein Complexes Exchange Rapidly in Living Drosophila.” Development 132 (17): 3963–76. https://doi.org/10.1242/dev.01950.

Fonseca, João Pedro, Philipp A. Steffen, Stefan Müller, James Lu, Anna Sawicka, Christian Seiser, and Leonie Ringrose. 2012. “In Vivo Polycomb Kinetics and Mitotic Chromatin Binding Distinguish Stem Cells from Differentiated Cells.” Genes & Development 26 (8): 857–71. https://doi.org/10.1101/gad.184648.111.

Francis, Nicole J., Robert E. Kingston, and Christopher L. Woodcock. 2004. “Chromatin Compaction by a Polycomb Group Protein Complex.” Science 306 (5701): 1574–77. https://doi.org/10.1126/science.1100576.

Franke, Axel, Sabine Messmer, and Renato Paro. 1995. “Mapping Functional Domains of the Polycomb Protein OfDrosophila Melanogaster.” Chromosome Research 3 (6): 351–60. https://doi.org/10.1007/BF00710016.

Grau, Daniel J., Brad A. Chapman, Joe D. Garlick, Mark Borowsky, Nicole J. Francis, and Robert E. Kingston. 2011. “Compaction of Chromatin by Diverse Polycomb Group Proteins Requires Localized Regions of High Charge.” Genes & Development 25 (20): 2210–21. https://doi.org/10.1101/gad.17288211.

Haddad, N., D. Jost, and C. Vaillant. 2017. “Perspectives: Using Polymer Modeling to Understand the Formation and Function of Nuclear Compartments.” Chromosome Research 25 (1): 35–50. https://doi.org/10.1007/s10577-016-9548-2.

Halverson, Jonathan D., Jan Smrek, Kurt Kremer, and Alexander Y. Grosberg. 2014. “From a Melt of Rings to Chromosome Territories: The Role of Topological Constraints in Genome Folding.” Reports on Progress in Physics 77 (2): 022601. https://doi.org/10.1088/0034-4885/77/2/022601.

Hou, Chunhui, Li Li, Zhaohui S. Qin, and Victor G. Corces. 2012. “Gene Density, Transcription, and Insulators Contribute to the Partition of the Drosophila Genome into Physical Domains.” Molecular Cell 48 (3): 471–84. https://doi.org/10.1016/j.molcel.2012.08.031.

Hug, Clemens B., Alexis G. Grimaldi, Kai Kruse, and Juan M. Vaquerizas. 2017. “Chromatin Architecture Emerges during Zygotic Genome Activation Independent of Transcription.” Cell 169 (2): 216–228.e19. https://doi.org/10.1016/j.cell.2017.03.024.

Huseyin, Miles K., and Robert J. Klose. 2021. “Live-Cell Single Particle Tracking of PRC1 Reveals a Highly Dynamic System with Low Target Site Occupancy.” Nature Communications 12 (1): 887. https://doi.org/10.1038/s41467-021-21130-6.

Ing-Simmons, Elizabeth, Roshan Vaid, Xin Yang Bing, Michael Levine, Mattias Mannervik, and Juan M. Vaquerizas. 2021. “Independence of Chromatin Conformation and Gene Regulation during Drosophila Dorsoventral Patterning.” Nature Genetics 53 (4): 487–99. https://doi.org/10.1038/s41588-021-00799-x.

Isono, Kyoichi, Takaho A. Endo, Manching Ku, Daisuke Yamada, Rie Suzuki, Jafar Sharif, Tomoyuki Ishikura, Tetsuro Toyoda, Bradley E. Bernstein, and Haruhiko Koseki. 2013. “SAM Domain Polymerization Links Subnuclear Clustering of PRC1 to Gene Silencing.” Developmental Cell 26 (6): 565–77. https://doi.org/10.1016/j.devcel.2013.08.016.

Jost, Daniel, Pascal Carrivain, Giacomo Cavalli, and Cédric Vaillant. 2014. “Modeling Epigenome Folding: Formation and Dynamics of Topologically Associated Chromatin Domains.” Nucleic Acids Research 42 (15): 9553–61. https://doi.org/10.1093/nar/gku698.

Kadoch, Cigall, Robert T. Williams, Joseph P. Calarco, Erik L. Miller, Christopher M. Weber, Simon M. G. Braun, John L. Pulice, Emma J. Chory, and Gerald R. Crabtree. 2017. “Dynamics of BAF–Polycomb Complex Opposition on Heterochromatin in Normal and Oncogenic States.” Nature Genetics 49 (2): 213–22. https://doi.org/10.1038/ng.3734.

Kaufman, Thomas C., Mark A. Seeger, and Gary Olsen. 1990. “Molecular and Genetic Organization of The Antennapedia Gene Complex of Drosophila Melanogaster.” In Advances in Genetics, edited by Theodore R. F. Wright, 27:309–62. Genetic Regulatory Hierarchies in Development. Academic Press. https://doi.org/10.1016/S0065-2660(08)60029-2.

Khokhlov, Alexei R., Alexander Yu Grosberg, and Vijay S. Pande. 1994. Statistical Physics of Macromolecules. Polymers and Complex Materials. AIP-Press. https://www.springer.com/gp/book/9781563960710.

Lanzuolo, Chiara, Virginie Roure, Job Dekker, Frédéric Bantignies, and Valerio Orlando. 2007. “Polycomb Response Elements Mediate the Formation of Chromosome Higher-Order Structures in the Bithorax Complex.” Nature Cell Biology 9 (10): 1167–74. https://doi.org/10.1038/ncb1637.

Leibler, Ludwik. 1980. “Theory of Microphase Separation in Block Copolymers.” Macromolecules 13 (6): 1602–17. https://doi.org/10.1021/ma60078a047.

Lewis, E. B. 1978. “A Gene Complex Controlling Segmentation in Drosophila.” Nature 276 (5688): 565–70. https://doi.org/10.1038/276565a0.

Lewis, Edward B., Barret D. Pfeiffer, David R. Mathog, and Susan E. Celniker. 2004. “Evolution of the Homeobox Complex in the Diptera.” In Genes, Development and Cancer: The Life and Work of Edward B. Lewis, edited by Howard D. Lipshitz, 381–85. Boston, MA: Springer US. https://doi.org/10.1007/978-1-4419-8981-9_24.

Li, Hua-Bing, Katsuhito Ohno, Hongxing Gui, and Vincenzo Pirrotta. 2013. “Insulators Target Active Genes to Transcription Factories and Polycomb-Repressed Genes to Polycomb Bodies.” PLOS Genetics 9 (4): e1003436. https://doi.org/10.1371/journal.pgen.1003436.

Lieberman-Aiden, Erez, Nynke L. van Berkum, Louise Williams, Maxim Imakaev, Tobias Ragoczy, Agnes Telling, Ido Amit, et al. 2009. “Comprehensive Mapping of Long-Range Interactions Reveals Folding Principles of the Human Genome.” Science 326 (5950): 289–93. https://doi.org/10.1126/science.1181369.

Linkert, Melissa, Curtis T. Rueden, Chris Allan, Jean-Marie Burel, Will Moore, Andrew Patterson, Brian Loranger, et al. 2010. “Metadata Matters: Access to Image Data in the Real World.” Journal of Cell Biology 189 (5): 777–82. https://doi.org/10.1083/jcb.201004104.

Liu, Miao, Yanfang Lu, Bing Yang, Yanbo Chen, Jonathan S.D. Radda, Mengwei Hu, Samuel G. Katz, and Siyuan Wang. 2020. “Multiplexed Imaging of Nucleome Architectures in Single Cells of Mammalian Tissue.” Nature Communications 11 (1). https://doi.org/10.1038/s41467-020-16732-5.

Loubiere, V., G. L. Papadopoulos, Q. Szabo, A-M. Martinez, and G. Cavalli. 2020. “Widespread Activation of Developmental Gene Expression Characterized by PRC1-Dependent Chromatin Looping.” Science Advances 6 (2): eaax4001. https://doi.org/10.1126/sciadv.aax4001.

Mateo, Leslie J., Sedona E. Murphy, Antonina Hafner, Isaac S. Cinquini, Carly A. Walker, and Alistair N. Boettiger. 2019. “Visualizing DNA Folding and RNA in Embryos at Single-Cell Resolution.” Nature 568 (7750): 49–54. https://doi.org/10.1038/s41586-019-1035-4.

Mirny, Leonid A. 2011. “The Fractal Globule as a Model of Chromatin Architecture in the Cell.” Chromosome Research 19 (1): 37–51. https://doi.org/10.1007/s10577-010-9177-0.

Nègre, Nicolas, Jérôme Hennetin, Ling V. Sun, Sergey Lavrov, Michel Bellis, Kevin P. White, and Giacomo Cavalli. 2006. “Chromosomal Distribution of PcG Proteins during Drosophila Development.” PLOS Biology 4 (6): e170. https://doi.org/10.1371/journal.pbio.0040170.

Nir, Guy, Irene Farabella, Cynthia Pérez Estrada, Carl G Ebeling, Brian J Beliveau, Hiroshi M Sasaki, S Dean Lee, et al. 2018. “Walking along Chromosomes with Super-Resolution Imaging, Contact Maps, and Integrative Modeling.” PLoS Genet. 14 (12): e1007872.

Nora, Elphège P., Bryan R. Lajoie, Edda G. Schulz, Luca Giorgetti, Ikuhiro Okamoto, Nicolas Servant, Tristan Piolot, et al. 2012. “Spatial Partitioning of the Regulatory Landscape of the X-Inactivation Centre.” Nature 485 (7398): 381–85. https://doi.org/10.1038/nature11049.

Nuebler, Johannes, Geoffrey Fudenberg, Maxim Imakaev, Nezar Abdennur, and Leonid A. Mirny. 2018. “Chromatin Organization by an Interplay of Loop Extrusion and Compartmental Segregation.” Proceedings of the National Academy of Sciences 115 (29): E6697–6706. https://doi.org/10.1073/pnas.1717730115.

Ogiyama, Yuki, Bernd Schuettengruber, Giorgio L. Papadopoulos, Jia-Ming Chang, and Giacomo Cavalli. 2018. “Polycomb-Dependent Chromatin Looping Contributes to Gene Silencing during Drosophila Development.” Molecular Cell 71 (1): 73–88.e5. https://doi.org/10.1016/j.molcel.2018.05.032.

Plys, Aaron J., Christopher P. Davis, Jongmin Kim, Gizem Rizki, Madeline M. Keenen, Sharon K. Marr, and Robert E. Kingston. 2019. “Phase Separation of Polycomb-Repressive Complex 1 Is Governed by a Charged Disordered Region of CBX2.” Genes & Development 33 (13–14): 799–813. https://doi.org/10.1101/gad.326488.119.

Rouillard, Jean-Marie, Michael Zuker, and Erdogan Gulari. 2003. “OligoArray 2.0: Design of Oligonucleotide Probes for DNA Microarrays Using a Thermodynamic Approach.” Nucleic Acids Research 31 (12): 3057–62. https://doi.org/10.1093/nar/gkg426.

Rowley, M. Jordan, Michael H. Nichols, Xiaowen Lyu, Masami Ando-Kuri, I. Sarahi M. Rivera, Karen Hermetz, Ping Wang, Yijun Ruan, and Victor G. Corces. 2017. “Evolutionarily Conserved Principles Predict 3D Chromatin Organization.” Molecular Cell 67 (5): 837–852.e7. https://doi.org/10.1016/j.molcel.2017.07.022.

Rubinstein, Michael, and Ralph H. Colby. 2003. Polymer Physics. Oxford, New York: Oxford University Press.

Sakaue, T., J.-C. Walter, E. Carlon, and C. Vanderzande. 2017. “Non-Markovian Dynamics of Reaction Coordinate in Polymer Folding.” Soft Matter 13 (17): 3174–81. https://doi.org/10.1039/C7SM00395A.

Saurin, Andrew J., Carol Shiels, Jill Williamson, David P.E. Satijn, Arie P. Otte, Denise Sheer, and Paul S. Freemont. 1998. “The Human Polycomb Group Complex Associates with Pericentromeric Heterochromatin to Form a Novel Nuclear Domain.” Journal of Cell Biology 142 (4): 887–98. https://doi.org/10.1083/jcb.142.4.887.

Schindelin, Johannes, Ignacio Arganda-Carreras, Erwin Frise, Verena Kaynig, Mark Longair, Tobias Pietzsch, Stephan Preibisch, et al. 2012. “Fiji: An Open-Source Platform for Biological-Image Analysis.” Nature Methods 9 (7): 676–82. https://doi.org/10.1038/nmeth.2019.

Schneider, Caroline A., Wayne S. Rasband, and Kevin W. Eliceiri. 2012. “NIH Image to ImageJ: 25 Years of Image Analysis.” Nature Methods 9 (7): 671–75. https://doi.org/10.1038/nmeth.2089.

Sexton, Tom, Eitan Yaffe, Ephraim Kenigsberg, Frédéric Bantignies, Benjamin Leblanc, Michael Hoichman, Hugues Parrinello, Amos Tanay, and Giacomo Cavalli. 2012. “Three-Dimensional Folding and Functional Organization Principles of the Drosophila Genome.” Cell 148 (3): 458–72. https://doi.org/10.1016/j.cell.2012.01.010.

Stanton, Benjamin Z., Courtney Hodges, Joseph P. Calarco, Simon M. G. Braun, Wai Lim Ku, Cigall Kadoch, Keji Zhao, and Gerald R. Crabtree. 2017. “Smarca4 ATPase Mutations Disrupt Direct Eviction of PRC1 from Chromatin.” Nature Genetics 49 (2): 282–88. https://doi.org/10.1038/ng.3735.

Steffen, Philipp A., João Pedro Fonseca, Cornelia Gänger, Eva Dworschak, Tobias Kockmann, Christian Beisel, and Leonie Ringrose. 2013. “Quantitative in Vivo Analysis of Chromatin Binding of Polycomb and Trithorax Group Proteins Reveals Retention of ASH1 on Mitotic Chromatin.” Nucleic Acids Research 41 (10): 5235–50. https://doi.org/10.1093/nar/gkt217.

Strübbe, Gero, Christian Popp, Alexander Schmidt, Andrea Pauli, Leonie Ringrose, Christian Beisel, and Renato Paro. 2011. “Polycomb Purification by in Vivo Biotinylation Tagging Reveals Cohesin and Trithorax Group Proteins as Interaction Partners.” Proceedings of the National Academy of Sciences 108 (14): 5572–77. https://doi.org/10.1073/pnas.1007916108.

Su, Jun-Han, Pu Zheng, Seon S. Kinrot, Bogdan Bintu, and Xiaowei Zhuang. 2020. “Genome-Scale Imaging of the 3D Organization and Transcriptional Activity of Chromatin.” Cell 182 (6): 1641–1659.e26. https://doi.org/10.1016/j.cell.2020.07.032.

Szabo, Quentin, Frédéric Bantignies, and Giacomo Cavalli. 2019. “Principles of Genome Folding into Topologically Associating Domains.” Science Advances 5 (4): eaaw1668. https://doi.org/10.1126/sciadv.aaw1668.

Szabo, Quentin, Daniel Jost, Jia-Ming Chang, Diego I. Cattoni, Giorgio L. Papadopoulos, Boyan Bonev, Tom Sexton, et al. 2018a. “TADs Are 3D Structural Units of Higher-Order Chromosome Organization in *Drosophila*.” Science Advances 4 (2): eaar8082. https://doi.org/10.1126/sciadv.aar8082.

Takei, Yodai, Jina Yun, Shiwei Zheng, Noah Ollikainen, Nico Pierson, Jonathan White, Sheel Shah, et al. 2021. “Integrated Spatial Genomics Reveals Global Architecture of Single Nuclei.” Nature 590 (7845): 344–50. https://doi.org/10.1038/s41586-020-03126-2.

Tolhuis, Bas, Marleen Blom, Ron M. Kerkhoven, Ludo Pagie, Hans Teunissen, Marja Nieuwland, Marieke Simonis, Wouter de Laat, Maarten van Lohuizen, and Bas van Steensel. 2011. “Interactions among Polycomb Domains Are Guided by Chromosome Architecture.” PLOS Genetics 7 (3): e1001343. https://doi.org/10.1371/journal.pgen.1001343.

Tuszynska, Irina, Pawel Bednarz, and Bartek Wilczynski. 2021. “Black Chromatin Is Indispensable for Accurate Simulations of Drosophila Melanogaster Chromatin Structure.” bioRxiv. https://doi.org/10.1101/2021.12.12.472204.

Ulianov, Sergey V., Vlada V. Zakharova, Aleksandra A. Galitsyna, Pavel I. Kos, Kirill E. Polovnikov, Ilya M. Flyamer, Elena A. Mikhaleva, et al. 2021. “Order and Stochasticity in the Folding of Individual Drosophila Genomes.” Nature Communications 12 (1): 41. https://doi.org/10.1038/s41467-020-20292-z.

Walter, J.-C., and G. T. Barkema. 2015. “An Introduction to Monte Carlo Methods.” Physica A: Statistical Mechanics and Its Applications, Proceedings of the 13th International Summer School on Fundamental Problems in Statistical Physics, 418 (January): 78–87. https://doi.org/10.1016/j.physa.2014.06.014.

Walter, J.-C., M. Baiesi, G. T. Barkema, and E. Carlon. 2013. “Unwinding Relaxation Dynamics of Polymers.” Physical Review Letters 110 (6): 068301. https://doi.org/10.1103/PhysRevLett.110.068301.

Wang, Liangjun, J. Lesley Brown, Ru Cao, Yi Zhang, Judith A Kassis, and Richard S Jones. 2004. “Hierarchical Recruitment of Polycomb Group Silencing Complexes.” Molecular Cell 14 (5): 637–46. https://doi.org/10.1016/j.molcel.2004.05.009.

Wani, Ajazul H., Alistair N. Boettiger, Patrick Schorderet, Ayla Ergun, Christine Münger, Ruslan I. Sadreyev, Xiaowei Zhuang, Robert E. Kingston, and Nicole J. Francis. 2016. “Chromatin Topology Is Coupled to Polycomb Group Protein Subnuclear Organization.” Nature Communications 7(1): 10291. https://doi.org/10.1038/ncomms10291.

Zenk, Fides, Yinxiu Zhan, Pavel Kos, Eva Löser, Nazerke Atinbayeva, Melanie Schächtle, Guido Tiana, Luca Giorgetti, and Nicola Iovino. 2021. “HP1 Drives de Novo 3D Genome Reorganization in Early Drosophila Embryos.” Nature 593 (7858): 289–93. https://doi.org/10.1038/s41586-021-03460-z.

