## Supplementary Data for "Multiplexed chromatin imaging reveals predominantly pairwise long-range coordination between *Drosophila* Polycomb genes"

**Index**

**Supplementary Figure 1**	**3**

Design of oligopaints library and quantification of Pc contacts from Hi-C data.	3

**Supplementary Figure 2**	**5**

Controls and supplementary data corresponding to Figure 1.	5

**Supplementary Figure 3**	**7**

Supplementary data corresponding to Figure 2.	7

**Supplementary Figure 4**	**9**

Expression patterns of Hox and non-Hox Pc genes from chr3R.	9

**Supplementary Figure 5**	**10**

Supplementary data corresponding to Figure 3.	10

**Supplementary Figure 6**	**11**

Supplementary data corresponding to Figure 4.	11

**Supplementary Table 1**	**12**

**Supplementary Table 2**	**14**

**Supplementary Table 3**	**17**

### Supplementary Figure 1

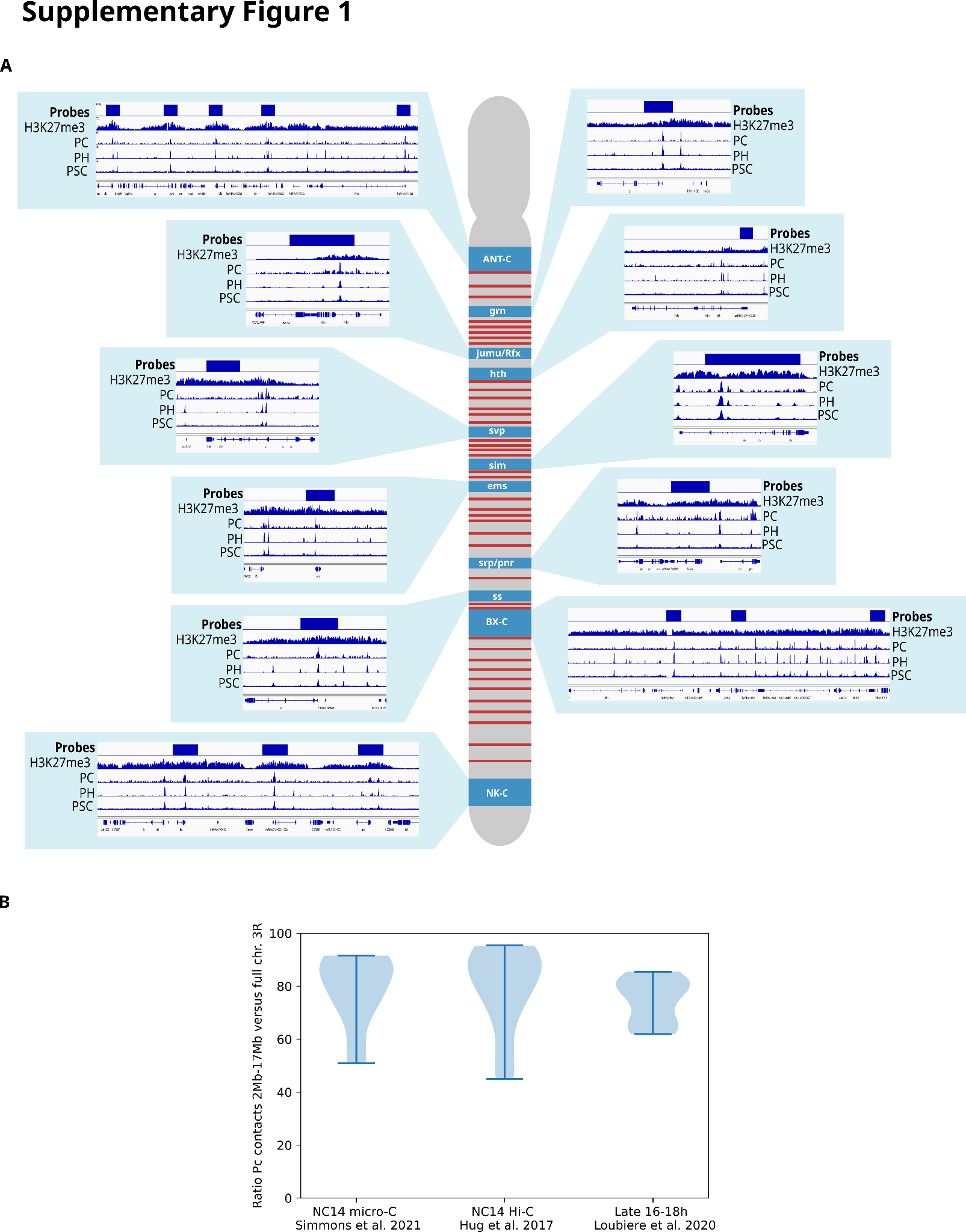

#### **Design of oligopaints library and quantification of Pc contacts from Hi-C data.**

**A.** Scheme representing chr. 3R and the Pc genomic loci used for this study (blue). The probe position within the locus is shown as a blue box, and is shown together with Chip-seq profiles for H3K27me3, PC, PH, and PSC. Regions displaying active chromatin regions (used in Fig. 2K) are shown in red.

**B.** Violin plots displaying the proportion of Pc target contacts in the selected ~15Mb portion of chr. 3R vs all Pc targets in chr. 3R, for several Hi-C and micro-C datasets (Ing-Simmons et al. 2021; Hug et al. 2017; Loubiere et al. 2020).

### Supplementary Figure 2

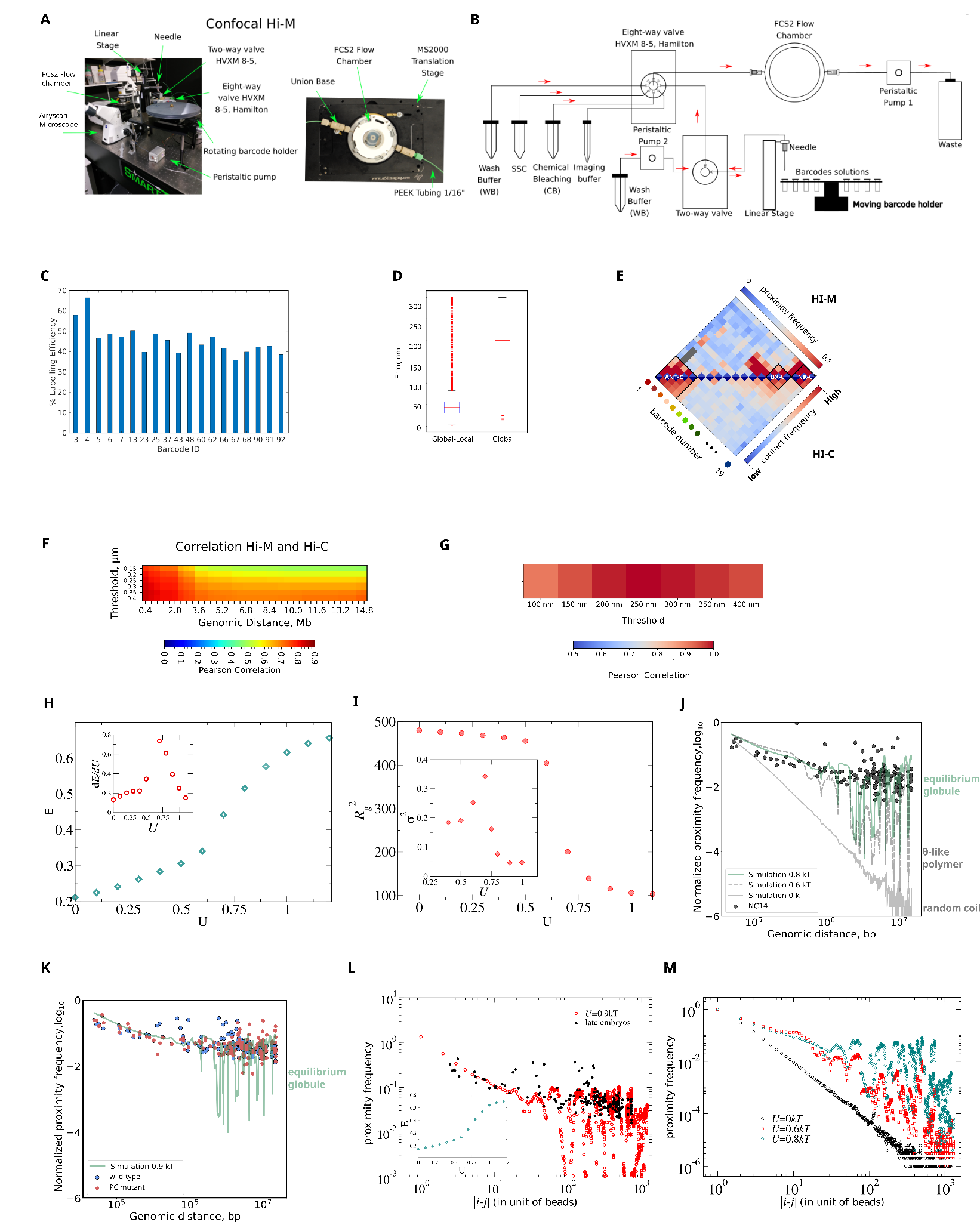

#### **Controls and supplementary data corresponding to Figure 1.**

**A - B.** Schematic representation of our confocal Hi-M microscope. Briefly, peristaltic pump 1 delivers buffers and barcodes into the microfluidics chamber. An 8-way valve selects the solution to be injected. A 2-way valve is used to either inject barcodes, or to rinse remaining barcode solutions in the needle between cycles, via peristaltic pump 2.

**C.** Labeling efficiency for all barcodes, defined as the percentage of nuclei displaying at least one detected barcode per imaging cycle.

**D.** Boxplot of residual error in drift correction of fiducial barcodes after global and local drift correction. Global correction is obtained by cross-correlating the fiducial images. Local correction is obtained as previously described (Cardozo Gizzi et al. 2020). The final correction vector is obtained by adding the global correction vector to the local correction vectors.

**E.** Comparison between Hi-M proximity frequency map and a Hi-C contact frequency map (Ogiyama et al. 2018) (wild-type S15-16 embryos).

**F.** Pearson correlation coefficients for different genomic distances between Hi-C contact frequency map (panel E) and Hi-M maps derived using different distance thresholds. For the threshold used in this work (T = 250 nm), the correlation between HiC and Hi-M was considerably high (>0.7) for all genomic distances.

**G.** Pearson correlation between the Hi-M proximity frequency map obtained with a distance threshold of T = 250 nm, and maps obtained using lower or higher distance thresholds.

**H.** Total energy (*E*) of the polymer (Fig. 1F, N=866 beads, 20kb / bead) as a function of the interaction energy between monomers (*U*). Inset represents the derivative of the energy (*dE/dU*) with a peak at the critical energy *U_c_*.

**I.** Radius of gyration (R_G_^2^ , in units of monomer length *a*) as a function of the interaction energy between monomers (*U*). Three different phases can be observed: coil regime (large values of R_G_), θ-regime (maximum in inset), and globule regime (small values of R_G_). Inset: fluctuations of R_G_^2^ as a function of U, defined as σ^2^ = (<R_G_^4^> - <R_G_^2^>^2^ )/ <R_G_^2^>^2^.

**J.** Normalized proximity frequency versus genomic distance for early NC14 embryos (gray circles) and from simulation of a self- interacting polymer in the globule (green curve, *U*=0.8k_B_T), θ-like (dashed gray) or random coil (solid gray) regimes. Experimental proximity frequency is normalized such that *P(s)* = 1 when the genomic distance *s* tends to 0.

**K.** Normalized proximity frequency versus genomic distance for wild-type (blue circles) and Pc mutant S15-S16 embryos (red circles), overlaid with that of a self-interacting polymer in the globule phase (green curve, *U* = 0.9k_B_T). Experimental proximity frequencies were normalized such that *P(s)* = 1 when the genomic distance *s* tends to *0*.

**L.** Experimental proximity frequencies versus genomic distance for wild-type S15-S16 embryos (black circles) and simulated proximity frequencies (red circles) for a lattice copolymer of the entire chr. 3R (N=1338 beads, 20kb / bead). Inset: Phase diagram (*E* versus *U*) for the entire chromosome chr. 3R. As in panel H, three different regimes corresponding to coil, θ and globule, can be readily observed.

**M.** Simulated proximity frequencies as a function of bead distance (*s*) for the entire chr. 3R, for different values of the interaction energy (*U*). As for the polymer in Fig. 1F, three main regimes can be observed: coil regime (black), θ regime (red), and globule regime (green).

### Supplementary Figure 3

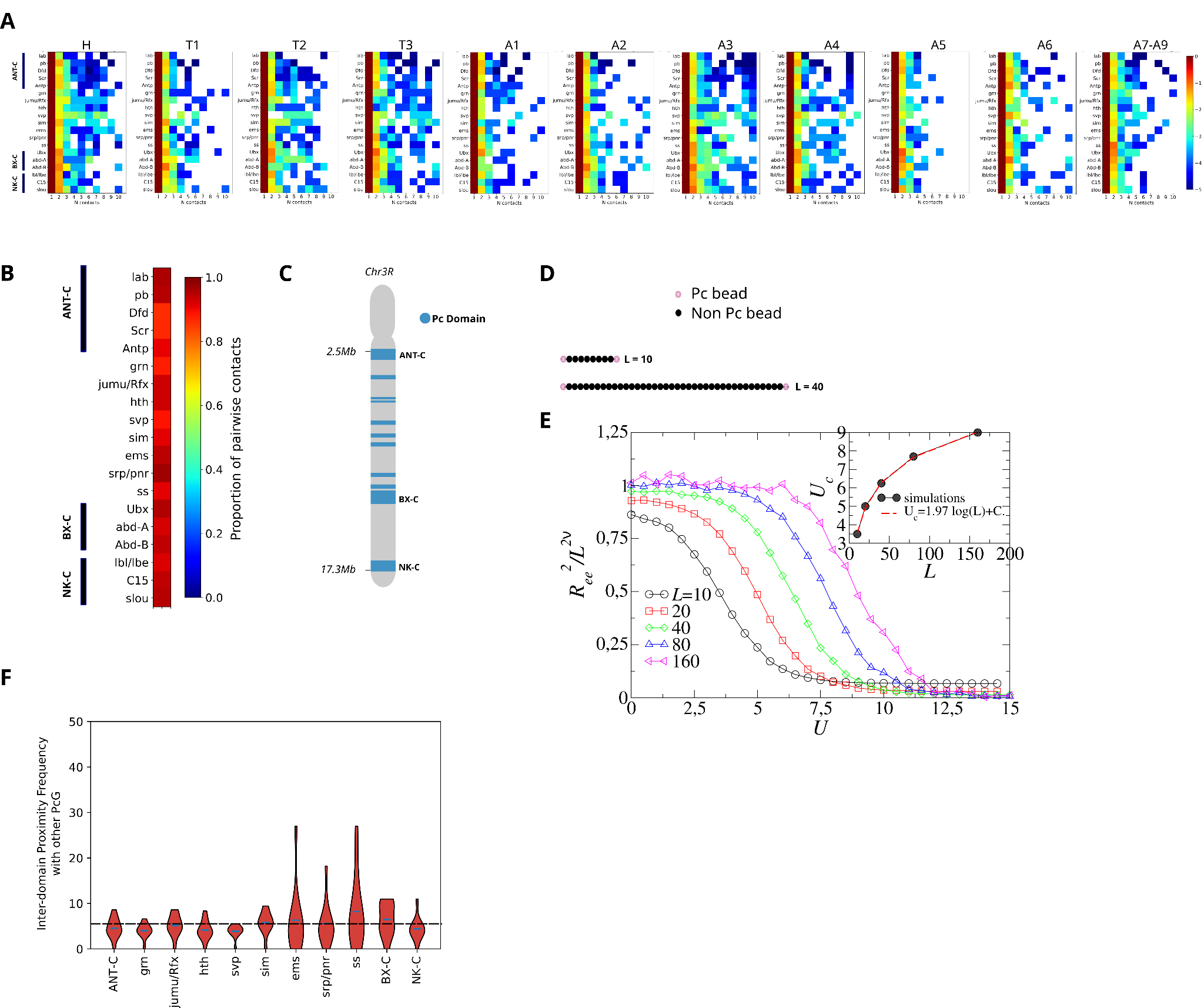

#### **Supplementary data corresponding to Figure 2.**

**A.** Frequency of multi-way clusters as a function of the number of targets in a cluster, normalized by the pairwise cluster frequency, for different segments of wild-type S15-S16 embryos. The frequency of multi-way interactions decreases with the number of co-localizing targets for all segments of the embryo, indicating that pairwise interactions are prevalent regardless of the tissue.

**B.** Frequencies of pairwise interactions among all multiway interactions for wild-type NC14 embryos.

**C.** Genomic distribution of Pc barcodes within a 1Mb window on chr. 3R. Barcodes closer than this genomic distance were fused together, as their physical distance is expected to be closer than the diffraction limit of light (see main text).

**D.** Cartoon representation of the toy model, where Pc beads are located at the ends of the chain of length L (L = 10 and L = 40 are shown as examples).

**E.** Rescaled end-to-end distance of the polymer as a function of the energy of interaction *U* between two attractively interacting beads separated by a polymeric chain of length L. The different curves correspond to different polymer lengths L. The energy needed to bridge the polymer increases with increasing polymer length (and thus increasing entropy). This illustrates that two distant Pc loci need to compete against the entropy of chromatin separating them, preventing the coalescence of several loci into phase-separated droplets. The rescaled end-to-end distance R_ee_/L^2𝜈^R_ee_/L^2𝜈^ of the polymer is proportional to the proximity frequency of the end beads, P_contact_ = 1 - R_ee_/L^2𝜈^. For small values of U, the polymer is in the coil regime, thus R_ee_/L^2𝜈^~1. As *U* increases, R_ee_/L^2𝜈^ decreases until the two Pc beads at the ends of the polymer stick together (for *U* >> *U_c_*). The critical energy (*U_c_*) increases logarithmically with the polymer length L as *U_c_*~3𝜈 log(L) (inset).

**F.** Violin plot distributions representing the normalized frequency with which Pc domains interact with other Pc domains in S15-S16 embryos. For each pair of barcodes, the normalized frequency was calculated as the ratio between the number of co-localized barcodes and the number of times those barcodes were detected. Grey line represents the mean. Dashed line represents the average normalized proximity frequency.

### Supplementary Figure 4

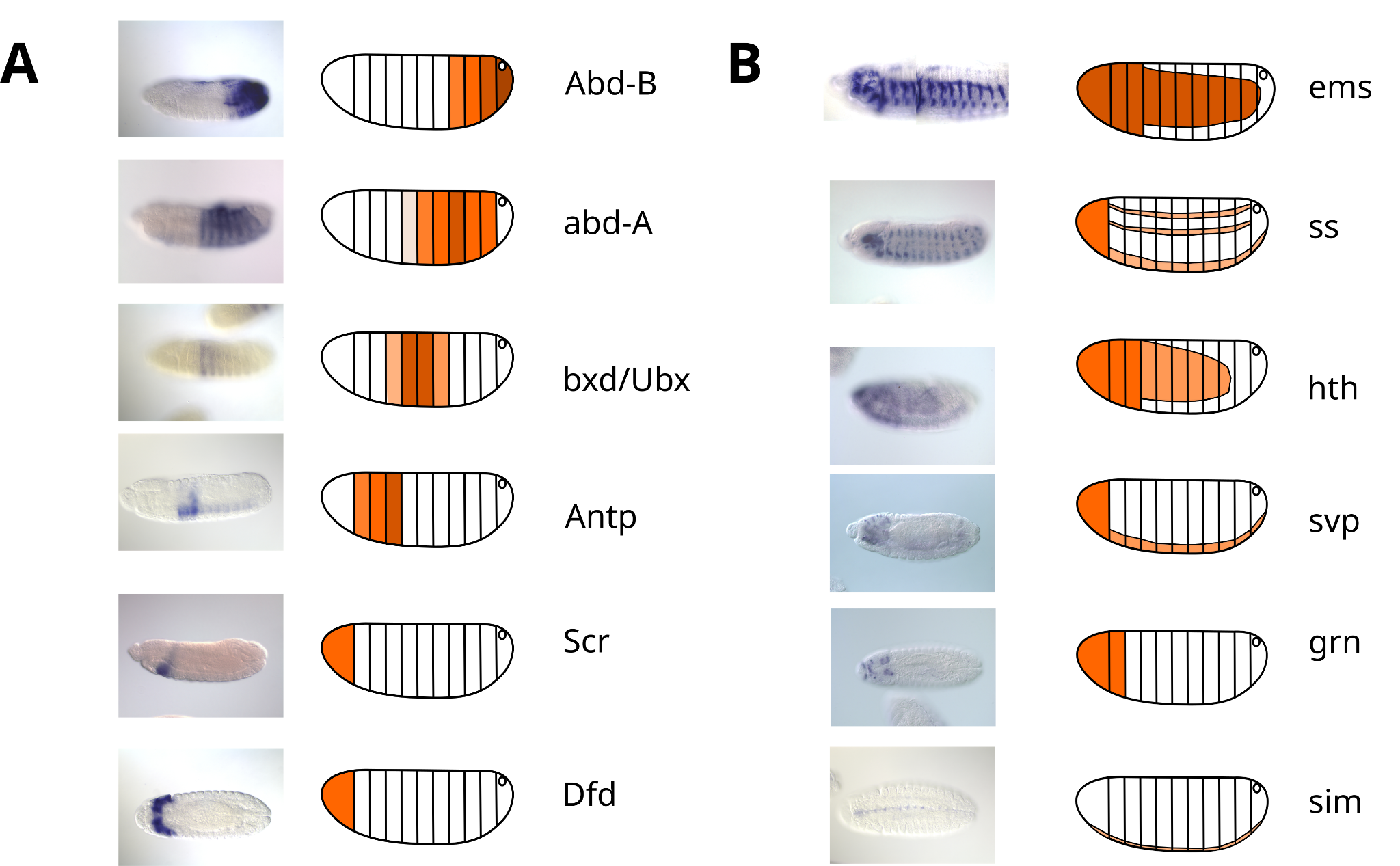

#### **Expression patterns of *Hox* and non-*Hox* Pc genes from chr3R.**

**A - B.** Expression patterns of Hox (A) and non-Hox (B) genes. Images are representative examples from the Berkeley database (<https://insitu.fruitfly.org/cgi-bin/ex/insitu.pl>).

### Supplementary Figure 5

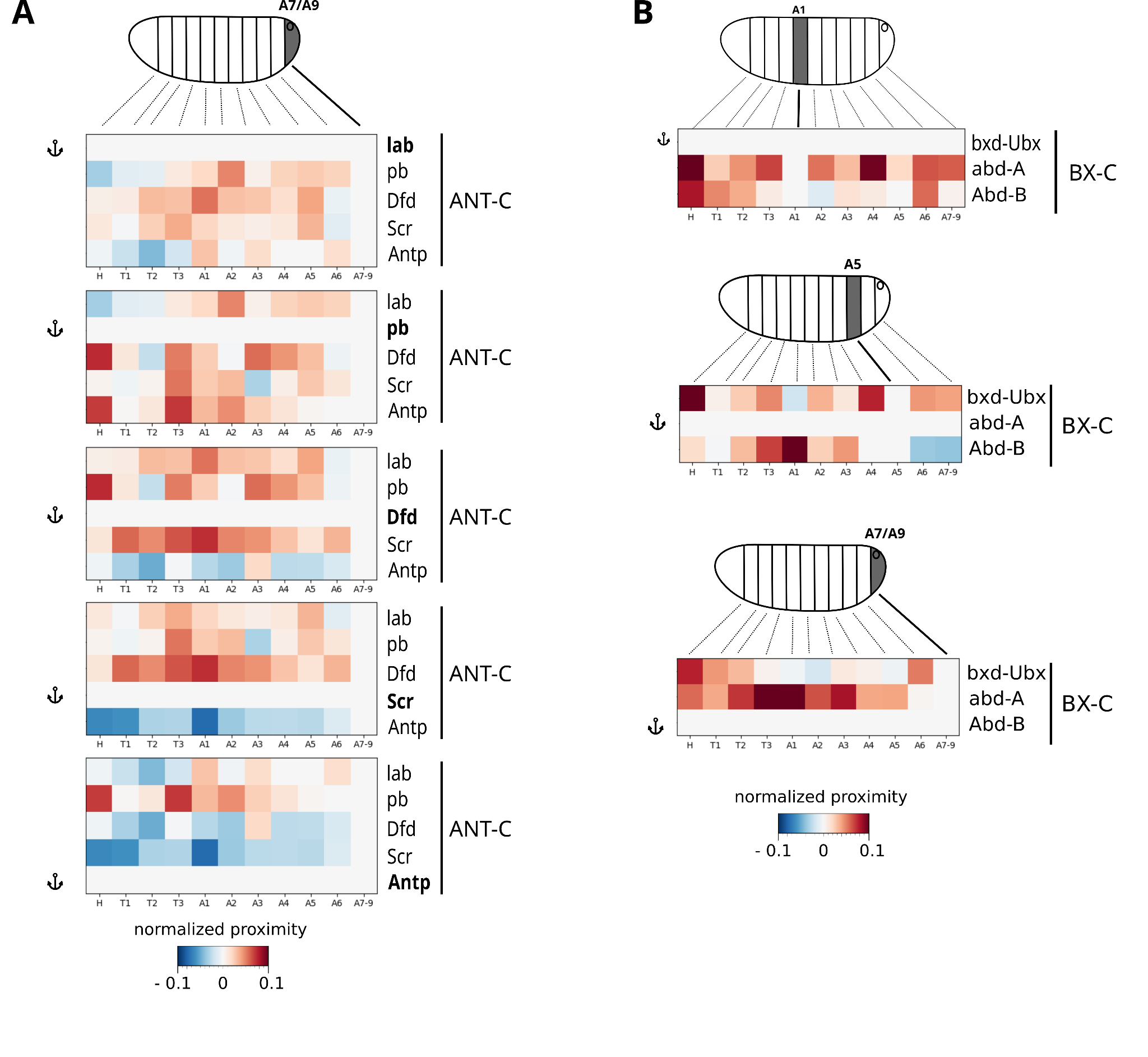

#### **Supplementary data corresponding to Figure 3.**

**A.** IntraTAD normalized proximity frequency maps for ANT-C gene targets, normalized in segments of repression. In most cases, the normalized proximity frequencies are positive, consistent with segregation of active from repressed targets. However, we note cases where the normalized proximity frequencies are negative, indicating a more complex picture.

**B.** IntraTAD normalized proximity frequency maps for BX-C gene targets (bxd/Ubx, Abd-a and Abd-B), normalized in their segments of expression. In most cases, normalized proximity frequencies are positive, indicating that BX-C targets segregate from other repressed targets within the TAD upon activation.

### Supplementary Figure 6

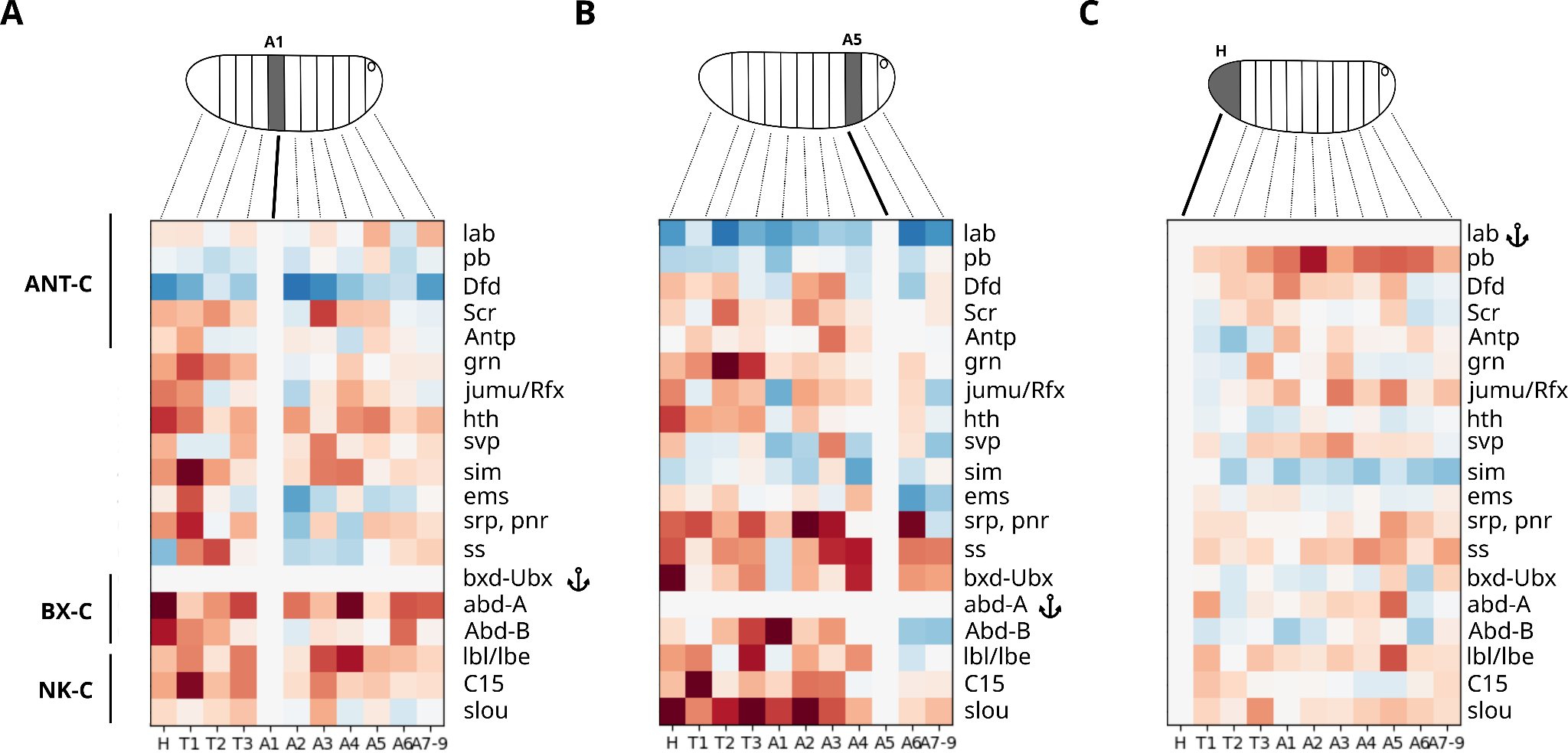

#### **Supplementary data corresponding to Figure 4.**

**A - C.** Normalized proximity frequency maps for a selection of anchors: bcd-Ubx (A), Abd-a (B) and Abd-B (C). Proximity frequencies were normalized by the segments of maximal expression of the anchor. In most cases, normalized proximity frequencies were positive, indicating that Hox PcG genes segregate from other PcG repressed targets upon activation.

### Supplementary Table 1

Sequences and IDs of Pc barcodes

| **Barcode ID** | **Sequence** |
| --- | --- |
| RT003-Imag_OligoRT1 | GCGCGATTGACCGTCTCGTTcaagtatgcaCACCGACGTCGCATAGAACGGAAGAGCGTGTG |
| RT004-Imag_OligoRT1 | CCGATGCGCAGCAATTCACTaagtcgtacgCACCGACGTCGCATAGAACGGAAGAGCGTGTG |
| RT005-Imag_OligoRT1 | GCCACGGTCCCGTTGAACTTcgaaacatcgCACCGACGTCGCATAGAACGGAAGAGCGTGTG |
| RT006-Imag_OligoRT1 | CGTCCAGCGCGTCAAACAGAacgaatccacCACCGACGTCGCATAGAACGGAAGAGCGTGTG |
| RT007-Imag_OligoRT1 | CCGTAACGAGCGTCCCTTGCcgcgaaatccCACCGACGTCGCATAGAACGGAAGAGCGTGTG |
| RT013-Imag_OligoRT1 | CGACGGATGTAATTCGGCCGgcctcgattaCACCGACGTCGCATAGAACGGAAGAGCGTGTG |
| RT023-Imag_OligoRT1 | TGGCGGTCTTAATATGCCCAgatggtcgacCACCGACGTCGCATAGAACGGAAGAGCGTGTG |
| RT025-Imag_OligoRT1 | ACCGAGCAGGTTAGTTGACGatgatccacgCACCGACGTCGCATAGAACGGAAGAGCGTGTG |
| RT037-Imag_OligoRT1 | CCGTATCCCTGGCGCGGACTcgtgcgggaaCACCGACGTCGCATAGAACGGAAGAGCGTGTG |
| RT043-Imag_OligoRT1 | ATCGGGCCCTTTTGTCTGACtaattccggtCACCGACGTCGCATAGAACGGAAGAGCGTGTG |
| RT048-Imag_OligoRT1 | GACGGCAAGAGAGCGTGCGTgccatggtacCACCGACGTCGCATAGAACGGAAGAGCGTGTG |
| RT060-Imag_OligoRT1 | GATGATCCGCTGAAGTCAAAgtgcgcgttaCACCGACGTCGCATAGAACGGAAGAGCGTGTG |
| RT062-Imag_OligoRT1 | AAGATGCCGTGAGCCTTTCAgcgcggtcccCACCGACGTCGCATAGAACGGAAGAGCGTGTG |
| RT066-Imag_OligoRT1 | CTATCGCTGATTCGTCCGGCgaaaccggcaCACCGACGTCGCATAGAACGGAAGAGCGTGTG |
| RT067-Imag_OligoRT1 | AGCCGCGAATCGGCCGTCTAatgcgatgagCACCGACGTCGCATAGAACGGAAGAGCGTGTG |
| RT068-Imag_OligoRT1 | AAATCCCGGAAATAACGGCCcgccttaccgCACCGACGTCGCATAGAACGGAAGAGCGTGTG |
| RT090-Imag_OligoRT1 | CCCTTGTGAGCGCCCGACATgcgttgatgtCACCGACGTCGCATAGAACGGAAGAGCGTGTG |
| RT091-Imag_OligoRT1 | CTCAAGCGGCAATCGTGTTGgttcgggcttCACCGACGTCGCATAGAACGGAAGAGCGTGTG |
| RT092-Imag_OligoRT1 | TGTCAGGTCCGCATGGGTCGcgcttatcgaCACCGACGTCGCATAGAACGGAAGAGCGTGTG |

### Supplementary Table 2

Sequences and IDs of active barcodes

| **Barcode ID** | **Sequence** |
| --- | --- |
| RT002-Imag_OligoRT1 | CCAGATCGGACGATCATGGGacaaatccgaCACCGACGTCGCATAGAACGGAAGAGCGTGTG |
| RT008-Imag_OligoRT1 | CCTGGCGTTGCGACGACTAAgcatgagttgCACCGACGTCGCATAGAACGGAAGAGCGTGTG |
| RT010-Imag_OligoRT1 | CCAGGTCCGTCACGCAATTTggccaatggcCACCGACGTCGCATAGAACGGAAGAGCGTGTG |
| RT012-Imag_OligoRT1 | CGCTTGTCGGGAACGGATACcgcgcggatcCACCGACGTCGCATAGAACGGAAGAGCGTGTG |
| RT014-Imag_OligoRT1 | CCGCTTGCGAGTAGGGCAATgcccgtattcCACCGACGTCGCATAGAACGGAAGAGCGTGTG |
| RT016-Imag_OligoRT1 | CCGTAAGGTGGGACCGAGGGagggtcatcgCACCGACGTCGCATAGAACGGAAGAGCGTGTG |
| RT018-Imag_OligoRT1 | ATCGCGCAATTGCGTAGTTCtgctagttcgCACCGACGTCGCATAGAACGGAAGAGCGTGTG |
| RT020-Imag_OligoRT1 | TTCCGCGTCATGTACCGGTTcgaccttaggCACCGACGTCGCATAGAACGGAAGAGCGTGTG |
| RT022-Imag_OligoRT1 | TGTCGCTACCGGCTTTGCTAgggaaacggtCACCGACGTCGCATAGAACGGAAGAGCGTGTG |
| RT026-Imag_OligoRT1 | ACGCTCGACAAGAACAGGACtagacgcaccCACCGACGTCGCATAGAACGGAAGAGCGTGTG |
| RT028-Imag_OligoRT1 | GTCTGGAACGACGGATTCAGgggtactcgcCACCGACGTCGCATAGAACGGAAGAGCGTGTG |
| RT030-Imag_OligoRT1 | CGTTCGTACCGCGTACTTCGagacgacgcaCACCGACGTCGCATAGAACGGAAGAGCGTGTG |
| RT032-Imag_OligoRT1 | TCGCCGCGTTTAGACGGGCTccaatgtaccCACCGACGTCGCATAGAACGGAAGAGCGTGTG |
| RT034-Imag_OligoRT1 | TACGGGCCCAGACGTTTCATgctacagcgtCACCGACGTCGCATAGAACGGAAGAGCGTGTG |
| RT036-Imag_OligoRT1 | CGATTAGCGCTCGTGCGCGAttaggtccggCACCGACGTCGCATAGAACGGAAGAGCGTGTG |
| RT038-Imag_OligoRT1 | ACGTAAGGCAGCTTGCGTTAaatccggcgtCACCGACGTCGCATAGAACGGAAGAGCGTGTG |
| RT040-Imag_OligoRT1 | TACGAGCCCTCTTGGACGGGgcgggattcgCACCGACGTCGCATAGAACGGAAGAGCGTGTG |
| RT042-Imag_OligoRT1 | CTCGGAGCGTTACTGCGGGCctttgttcggCACCGACGTCGCATAGAACGGAAGAGCGTGTG |
| RT045-Imag_OligoRT1 | TGACTCGGCTCAGTCGCGGCatcggaacggCACCGACGTCGCATAGAACGGAAGAGCGTGTG |
| RT047-Imag_OligoRT1 | CGGCGTCGGTAGGCCTTCGCttacgaaggtCACCGACGTCGCATAGAACGGAAGAGCGTGTG |
| RT049-Imag_OligoRT1 | GTCCCGGAAGCACGGGCGACcggattggtcCACCGACGTCGCATAGAACGGAAGAGCGTGTG |
| RT051-Imag_OligoRT1 | CGTCGTGGGACGTGGACCGTacacccgataCACCGACGTCGCATAGAACGGAAGAGCGTGTG |
| RT053-Imag_OligoRT1 | ATCGCCTCAATAAAGGCGACctttacgcggCACCGACGTCGCATAGAACGGAAGAGCGTGTG |
| RT055-Imag_OligoRT1 | CGTAGCTCACCCGGTTAGCGgatcgaccggCACCGACGTCGCATAGAACGGAAGAGCGTGTG |
| RT057-Imag_OligoRT1 | ACTCTATCGGCATGGAGGTAgggcgacgggCACCGACGTCGCATAGAACGGAAGAGCGTGTG |
| RT059-Imag_OligoRT1 | GACAATGGTGCGAAAGACCCgccatgcgtcCACCGACGTCGCATAGAACGGAAGAGCGTGTG |
| RT061-Imag_OligoRT1 | CGCGAGGAATTGGCGCATCGttgagcgttgCACCGACGTCGCATAGAACGGAAGAGCGTGTG |
| RT063-Imag_OligoRT1 | CTTCCAATCCCTAAGGCCACaccgaatcggCACCGACGTCGCATAGAACGGAAGAGCGTGTG |
| RT065-Imag_OligoRT1 | AACTTAGCGATCACGCGCAAcggatcgcccCACCGACGTCGCATAGAACGGAAGAGCGTGTG |
| RT069-Imag_OligoRT1 | AATGGGCATTCGATGCGCAGgcaacgggcaCACCGACGTCGCATAGAACGGAAGAGCGTGTG |
| RT071-Imag_OligoRT1 | CCAATGAAGCAACGCGCTTTcagcaggcgtCACCGACGTCGCATAGAACGGAAGAGCGTGTG |
| RT073-Imag_OligoRT1 | CTTTGTCGCGCCTTACACCAtgcgggtgcgCACCGACGTCGCATAGAACGGAAGAGCGTGTG |
| RT075-Imag_OligoRT1 | GAGAAGCGCTCGGGTATGACgcgtcgatcgCACCGACGTCGCATAGAACGGAAGAGCGTGTG |
| RT077-Imag_OligoRT1 | TGAAAGCCGGACAGTTCGCAtcggtcaaccCACCGACGTCGCATAGAACGGAAGAGCGTGTG |
| RT079-Imag_OligoRT1 | CACGATAGGCCCATCGCGAAgttgttcacgCACCGACGTCGCATAGAACGGAAGAGCGTGTG |
| RT081-Imag_OligoRT1 | TCGGAGTCAGCCTCGCGACTtcgcttgtgtCACCGACGTCGCATAGAACGGAAGAGCGTGTG |
| RT083-Imag_OligoRT1 | ACGCGTCGCCACACGCAATGgttcctctcaCACCGACGTCGCATAGAACGGAAGAGCGTGTG |
| RT084-Imag_OligoRT1 | CTTCACCGCGTTTGTTCAACcggtgccaagCACCGACGTCGCATAGAACGGAAGAGCGTGTG |
| RT086-Imag_OligoRT1 | GACACGTCCCTCAATCGAACtggccgacgcCACCGACGTCGCATAGAACGGAAGAGCGTGTG |
| RT088-Imag_OligoRT1 | TGGACTTCCGGTCGATATCCgggcgggagtCACCGACGTCGCATAGAACGGAAGAGCGTGTG |

### Supplementary Table 3

Sequence and IDs of adaptor, displacement, imaging, and fiducial oligos.

| **ID** | **Sequence** |
| --- | --- |
| Imaging oligo: MER1-SS-A647-32pb | CACACGCTCTTCCGTTCTATGCGACGTCGGTG/iThioMC6-D//3AlexF647N/ |
| Fiducial imaging oligo: MER23-RhoRed | CATTGCCGTATGGGCTAGGATGACCTGGCTCG/3RhodRd-XN/ |
| Fiducial adaptor oligo: RT008-Imag_OligoRT23 | CCT GGC GTT GCG ACG ACT AAG CAT GAG TTG CGA GCC AGG TCA TCC TAG CCC ATA CGG CAA TG |
| Fiducial displacement oligo: Displmt_RT008 | caactcatgcTTAGTCGTCGCAACGCCAGG |
